# Cryo-EM structures of the CDK11-cyclin L-SAP30BP complex reveal mechanisms of CDK11 regulation

**DOI:** 10.64898/2026.03.24.713564

**Authors:** Amy J. S. McGeoch, Victoria I. Cushing, Theodoros I. Roumeliotis, Nora B. Cronin, Stephen J. Hearnshaw, Jyoti S. Choudhary, Claudio Alfieri, Basil J. Greber

**Author notes:** correspondence to (B.J.G.).

## Abstract

The cyclin-dependent kinase CDK11 functions in transcription, mitotic progression, and mRNA splicing. Specifically, spliceosome activation during the B to B^act^ transition depends on phosphorylation of the U2 snRNP component SF3B1 by the CDK11-cyclin L-SAP30BP complex. Here, we present the structure of this spliceosome-activating CDK-cyclin complex, determined by cryogenic electron microscopy at 2.3 Å resolution. Our structure and biochemical experiments show that SAP30BP forms extensive interactions with cyclin L2, thereby stabilising it, and forms critical interactions with the C-terminal kinase lobe of CDK11 that promote complex assembly. Destabilisation of cyclin L2 in the absence of SAP30BP suggests that these principles are applicable to all CDK11-cyclin L complexes. Furthermore, we identify a pseudo-substrate sequence near the CDK11 C-terminus and provide evidence for a role of this segment in CDK11 auto-regulation. Finally, the structure of the CDK11-cyclin L2-SAP30BP complex bound to the clinical high-affinity CDK11 inhibitor OTS964 and a comparison to OTS964-bound off-target complexes provide insight into the mechanism of OTS964 selectivity and specificity.

## INTRODUCTION

The cyclin-dependent protein kinase (CDK) family comprises 20 members in human cells, most of which form binary complexes with one or several members of the cyclin family ^1^. CDKs play critical roles in cell growth and proliferation by controlling multiple key cellular pathways. The canonical role of CDK-cyclin complexes is in cell cycle control, where the enzymatic activities of CDKs oscillate due to a complex interplay of regulatory mechanisms, including cyclin availability ^2^. Additional CDKs that are usually constitutively associated with their partner cyclins perform prominent roles in transcription control and mRNA splicing ^1^. CDK11 is a representative of this latter group.

CDK11 is expressed in several variants and isoforms in human cells. Human CDK11 is encoded by two genes, and the resulting full-length proteins, CDK11A^p110^ and CDK11B ^p110^, are 97% identical on the sequence level ^1^. Further CDK11 diversity originates from alternatively spliced isoforms ^3^, and from the expression of a shorter CDK11 variant called CDK11^p58^, which arises from translation initiation at a cell-cycle-regulated internal ribosomal entry site ^4^. Therefore, the N-terminus of the resulting protein is truncated compared to the full-length CDK11^p110^ isoform (Supplementary Fig. 1a) ^3–5^. The CDK11^p58^ isoform is expressed in a cell-cycle specific manner and is required for progression through mitosis ^6,7^ while CDK11^p110^ is constitutively expressed and is involved in transcription and splicing ^8–12^. Both CDK11^p110^ and CDK11^p58^ form stable complexes with cyclins L1 and L2, which are encoded by distinct genes ^13–15^.

Early work on the involvement of CDK11 in splicing documented its association with splicing factors such as splicing regulator (SR) proteins ^8,13,15^. However, a mechanistic understanding of the role of CDK11 in the core splicing pathway remained elusive until the recent discovery that CDK11 inhibition impairs phosphorylation of the protein SF3B1, a key component of the SF3B complex within the U2 small nuclear ribonucleoprotein (U2 snRNP). The resulting deficiency in SF3B1 phosphorylation prevents spliceosomal activation, thereby stalling the spliceosome between the B- and B^act^-complex stages ^16^. This causes widespread intron retention ^16,17^, revealing CDK11 as a key player in spliceosome activation. This critical activity of CDK11 requires the formation of a trimeric spliceosome-activating CDK-cyclin complex that is composed of CDK11^p110^, cyclin L1/2, and SAP30BP ^17^. SAP30BP is a transcription and splicing factor that has been implicated in splicing of short introns ^18,19^. In the context of the CDK11-bound complex, SAP30BP stabilizes the CDK11-cyclin L complex, contributes to CDK11 activation, and mediates interactions with spliceosomal components ^17^.

Due to their important roles in cell growth and proliferation, CDKs have been the target of intense drug discovery efforts, with a focus on CDKs 4, 6, 7, and 9 ^20,21^. In addition to these intensively studied CDKs, CDK11 is an emerging drug target ^22,23^ because its activity is required for the growth of multiple cancer cell lines ^24,25^. Even though dedicated inhibitors for CDK11 have been discovered ^26^, one of the clinically most advanced CDK11 inhibitors, OTS964, was originally mischaracterised as a PBK (TOPK) inhibitor ^27^ and only later discovered to potently inhibit CDK11 ^28^. This inhibitor is currently in clinical evaluation for its activity against CDK11, and it has been instrumental in characterising the role of CDK11 in splicing ^16^.

Despite the critical importance of CDK11 and its cyclin L and SAP30BP partners for our understanding of fundamental biological processes and for cancer drug discovery, structures of full trimeric CDK11-cyclin L-SAP30BP complexes have remained elusive. This hampers our understanding of splicing activation and CDK11 regulation. Here, we report the structure of the fully assembled CDK11B-cyclin L2-SAP30BP trimer in complex with AMP-PNP or the inhibitor OTS964 along with structures of OTS964 bound to the off-targets CDK2-cyclin A2 and CDK7-cyclin H-MAT (the CDK-activating kinase, or CAK). Based on these results, we test structure-derived hypotheses on complex assembly and CDK11 regulation by biochemical experiments. Our work provides insight into the architecture and assembly of the activated CDK11 complex, reveals an auto-regulatory pseudo-substrate segment near the CDK11 C-terminus, and contributes to our understanding of the molecular basis of OTS964 selectivity.

## RESULTS AND DISCUSSION

### Structure determination

To investigate the mechanism by which SAP30BP promotes CDK11-cyclin L complex formation and CDK11 activation, we recombinantly expressed the trimeric CDK11-cyclin L-SAP30BP assembly in insect cells and purified the complex by affinity and gel filtration chromatography (Supplementary Fig. 1b-d). To facilitate structure determination, we truncated the N-terminus of CDK11, generating CDK11^p58^ (CDK11B^p110^ residues 357-795) and removed the C-terminal tail of cyclin L2, including its predicted disordered RS-domain, resulting in expression of cyclin L2 residues 1-319 (Supplementary Fig. 1a, b). Attempts to prepare CDK11-cyclin L2 complexes or express cyclin L2 on its own failed (Fig. 1a). This is consistent with previous reports indicating instability of cyclin L in the absence of SAP30BP, both when recombinantly expressed ^29^ and in the native context in human cells ^17^. These findings indicate that SAP30BP is an obligate binding partner of cyclin L and suggest that SAP30BP is likely to be a component of both CDK11^p110^ and CDK11^p58^-containing complexes.

**Figure 1.**
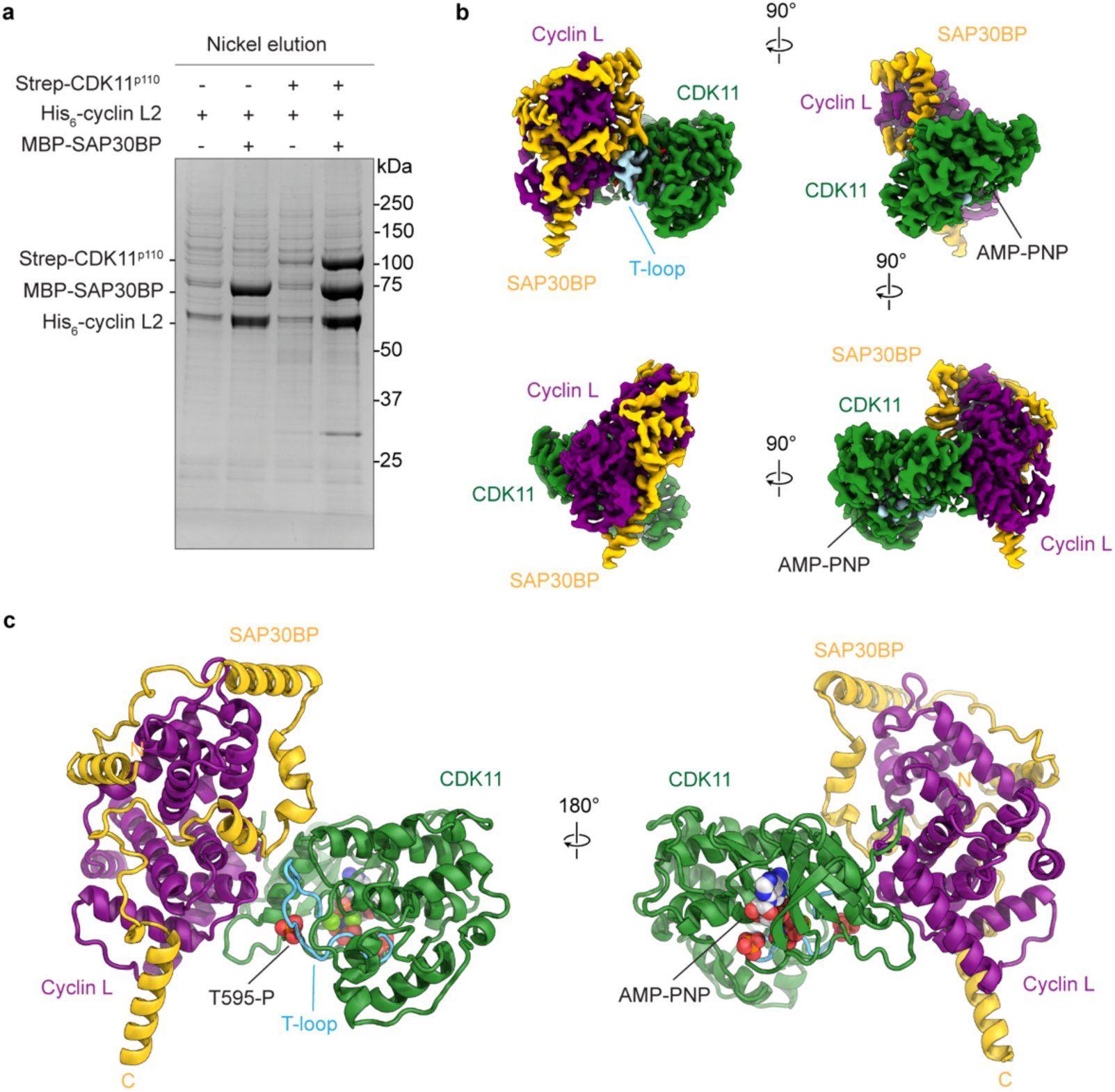
Reconstitution and cryo-EM structure of the CDK11-cyclin L-SAP30BP complex. (**a**) Result of small-scale Nickel-sepharose pulldown of complexes after co-expression of the indicated subunits in insect cells. In the absence of SAP30BP, cyclin L2 recovery is strongly diminished. (**b**) Cryo-EM map of the CDK11-cyclin L-SAP30BP complex in four orientations (related by 90° rotations) and with subunits distinctly coloured (CDK11 green, T-loop blue, cyclin L purple, SAP30BP yellow, AMP-PNP white). (**c**) Atomic model derived from the cryo-EM map shown in b. The subunits are coloured as in b; additionally, the phosphorylated T-loop residue T595 and the N- and C-terminal ends of the modelled portion of SAP30BP are indicated.

Using this reconstituted complex, we obtained a 2.3 Å-resolution cryo-EM reconstruction of the core module of the CDK11-cyclin L2-SAP30BP complex, visualising approx. 76 kDa in molecular weight (Fig. 1b, Supplementary Fig. 2a-d). The excellent quality of our cryo-EM map enabled building and refinement of a full atomic model and a detailed analysis of molecular contacts (Supplementary Fig. 2e-g). It reveals the molecular architecture of the CDK11-cyclin L2-SAP30BP complex, its interactions with the nucleotide analogue AMP-PNP (Supplementary Fig. 2e), and multiple phosphorylated residues on CDK11 (Fig. 1c, Supplementary Fig. 2g).

Structural modelling using AlphaFold3 ^30^ suggests that the highly charged N-terminal extension of CDK11^p110^, much of which consists of low-complexity regions (Supplementary Fig. 1a), is disordered in solution and does not contribute to folding and assembly of the core region of the CDK11-cyclin L2-SAP30BP complex (Supplementary Fig. 3). Therefore, the parts of CDK11^p110^ that are absent in our cryo-EM specimen are likely disordered in the free trimeric complex, suggesting that our structural analysis is valid for both CDK11^p58^ and CDK11^p110^ in the context of CDK11-cyclin L-SAP30BP complexes. Similarly, the sequence identity between cyclins L1 and L2 within the region observed in our structure is high (82% for residues 67-302), and the general conclusions are valid for both cyclin L variants. We will therefore use the designations CDK11 and cyclin L unless we refer to a certain variant of these proteins specifically.

### Architecture of the CDK11-cyclin L-SAP30BP complex and mechanism of its stabilisation by SAP30BP

Our structure shows that CDK11 and cyclin L form a canonical CDK-cyclin pair in which direct interactions are primarily mediated by the N-terminal kinase lobe of CDK11 and the N-terminal cyclin fold of cyclin L (Fig. 2a, b). The T-loop of CDK11 is found in an extended, active conformation, and the features of our cryo-EM map indicate that it is phosphorylated at T595 (CDK11B^p110^ residue numbering; Fig. 1c, Supplementary Fig. 2g). The phosphate moiety is coordinated by three arginine residues (R484, R585, R561), which thereby stabilise the active T-loop conformation, a feature that is commonly found in CDKs (Fig. 2b, Supplementary Fig. 2g).

**Figure 2.**
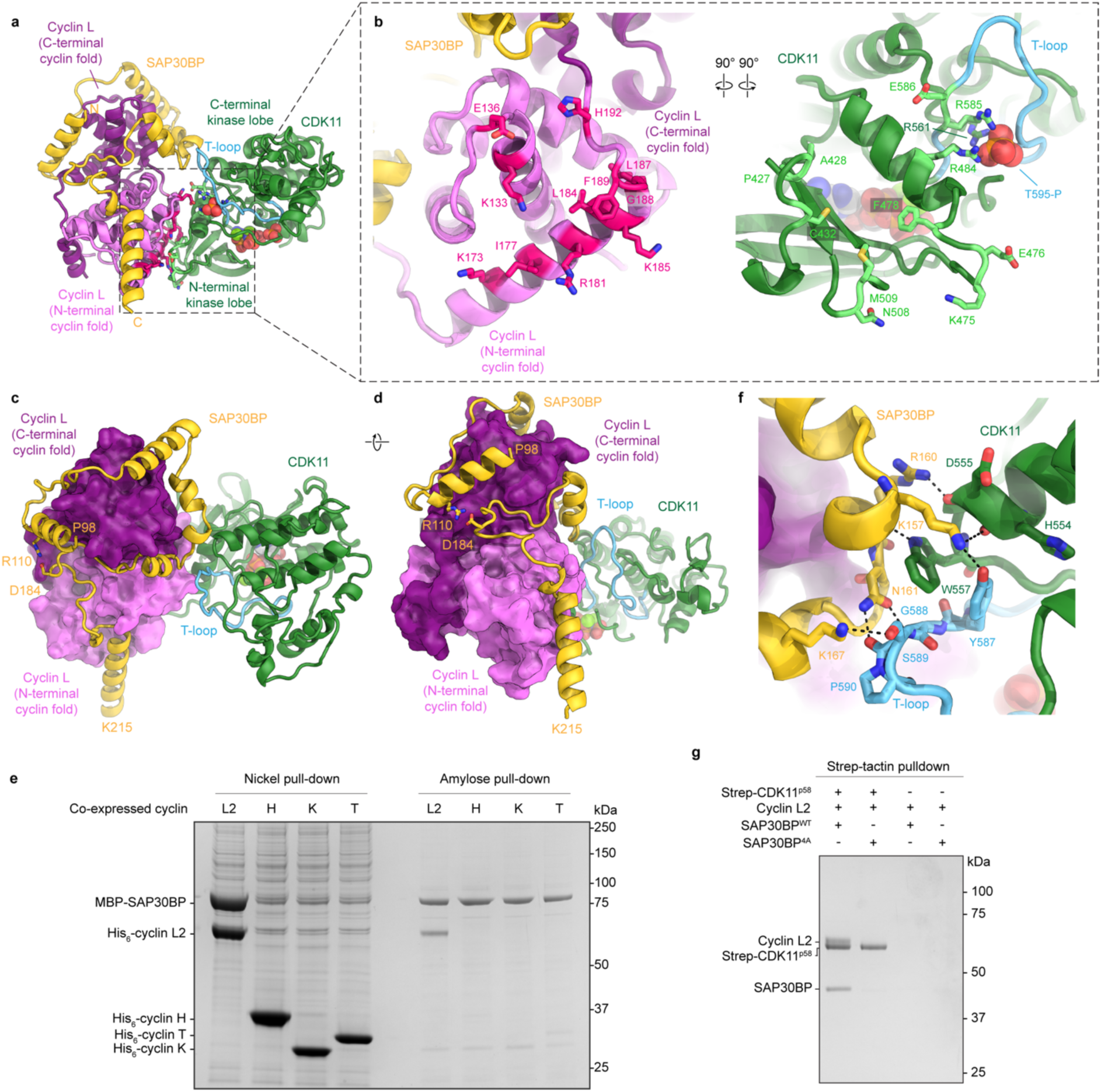
Molecular basis of CDK11-cyclin L-SAP30BP complex assembly. (**a**, **b**) Interactions between CDK11 and cyclin L. CDK11 is coloured green, with residues forming the interface with cyclin L coloured light green (distance cut-off: 3.6 Å) and the T-loop in blue; cyclin L is coloured pink and purple (N- and C-terminal cyclin folds, respectively), with residues forming the interface with CDK11 coloured bright red. SAP30BP is coloured yellow, and the N- and C-terminal ends of the model are labelled. (**c**, **d**) Interactions of SAP30BP with cyclin L (colours as in a-c). (**e**) Small-scale co-purification assay from insect cell lysates co-expressing MBP-SAP30BP with His_6_-tagged cyclins L2, H, K, and T. Only His_6_-cyclin L2 co-purifies MBP-SAP30BP (nickel-sepharose elution), and MBP-SAP30BP only co-purifies His_6_-cyclin L2 (amylose elution). (**f**) Interactions of SAP30BP with CDK11 (colours as in a). Hydrogen bonds are indicated as black dashed lines. (**g**) *In vitro* pulldown of purified cyclin L2 in complex with SAP30BP wild type (WT) and 4A mutant. Immobilised Strep-CDK11^p58^ retains cyclin L2-SAP30BP^WT^ but not cyclin L2-SAP30BP^4A^. The right-hand two lanes are negative controls without pre-incubation of the resin with Strep-CDK11^p58^.

Our cryo-EM map visualises a SAP30BP segment comprising residues from P98 to K215, which predominantly interacts with cyclin L (Fig. 2c, d). The C-terminal U2AF ligand motif (ULM) of SAP30BP, which engages in an interaction with the U2AF homology motif (UHM) of the splicing factor RBM17 to chaperone RBM17 to phosphorylated SF3B1 (ref. ^19^), is not visualised in our cryo-EM map. Therefore, this binding site for RBM17 on SAP30BP likely remains solvent-exposed and accessible to other proteins in the CDK-cyclin-bound complex. This may allow targeting of the entire trimeric complex to spliceosomal sites of action. Similarly, SAP30BP residues 73-88 encoded by its alternatively spliced exon 3 lie outside the region interacting with CDK11 and cyclin L, making this region available to form isoform-dependent interactions with mRNA biogenesis and processing factors, thereby regulating transcription and the alternative splicing of short introns ^18^.

SAP30BP residues 98-184 form six α-helical segments connected by extended linkers that encircle the C-terminal cyclin fold of cyclin L. This ring around cyclin L is sealed by a salt bridge between SAP30BP residues R110 and D184 (Fig. 2c, d). Subsequent residues of SAP30BP interact with the N-terminal cyclin fold of cyclin L, terminating in a long α-helix that extends into the solvent and likely continues beyond the last residue (K215) modelled in our structure (Fig. 1c, 2c, d). These extensive protein-protein interactions with both cyclin domains bury approx. 3530 Å^2^ in surface area ^31^, rationalising why cyclin L stability is compromised in the absence of SAP30BP (Fig. 1a and refs ^17,29^). Tight binding and formation of a stable complex between cyclin L and SAP30BP are in agreement with previous biochemical analysis ^17^. These interactions appear to be highly specific to cyclin L, as SAP30BP did not form stable complexes when co-expressed with the phylogenetically related cyclins H, K, and T (Fig. 2e) ^32^.

In contrast to the extensive interactions between cyclin L and SAP30BP, the molecular interface between SAP30BP and CDK11 are rather modest in extent; it is limited to an 11-residue segment of SAP30BP (residues 157-167) that contacts the C-terminal kinase lobe of CDK11 and residues Y587 and S589 in the CKD11 T-loop (Fig. 2f, Supplementary Fig. 2f). These interactions bury approximately 360 Å^2^ of surface area ^31^. They comprise multiple side chain-side chain and side chain-main chain hydrogen bonds and a cation-π interaction between CDK11 W557 and SAP30BP K157. Engineered mutations in the SAP30BP residues responsible for forming these interactions (K157A, R160A, N161A, K167A; SAP30BP^4A^) reduce the stability of the interaction of cyclin L-SAP30BP complexes with CDK11 (Fig. 2g). The contacts between SAP30BP and the CDK11 T-loop may be subject to modulation by post-translational modifications. Experiments using *Drosophila* CDK11 indicate that phosphorylation of S589 (human numbering; S712 in *Drosophila*) may play a regulatory role by suppressing CDK11 activity ^33^. In our cryo-EM map, this residue is un-phosphorylated and forms a charged hydrogen bond with SAP30BP K167 (Fig. 2f, Supplementary Fig. 2f).

Our structure thus provides a structural view of how SAP30BP promotes both cyclin L stability and CDK11-cyclin L complex formation. These functions of SAP30BP have been shown to critically contribute to CDK11 activity in splicing ^17^ and are likely to also support the other cellular functions of CDK11.

### CDK11 contains an auto-regulatory pseudo-substrate segment that can occlude the substrate binding site

The CDK11 active site in our cryo-EM structure is found in a state previously observed in complexes primed for phosphoryl transfer ^34^, in which the phosphates of the bound AMP-PNP molecule are coordinated to two magnesium ions and the catalytic lysine (K467 in CDK11^p110^) (Fig. 3a). The base of the AMP-PNP molecule is bound to backbone atoms of N517 and V519 in the hinge region of CDK11 (Fig. 3a, Supplementary Fig. 2e), as is typically observed in kinase structures ^35^, and its exocyclic amino group is within hydrogen bonding distance of the sulphur atom of the gatekeeper residue M516 (Fig. 3a).

**Figure 3.**
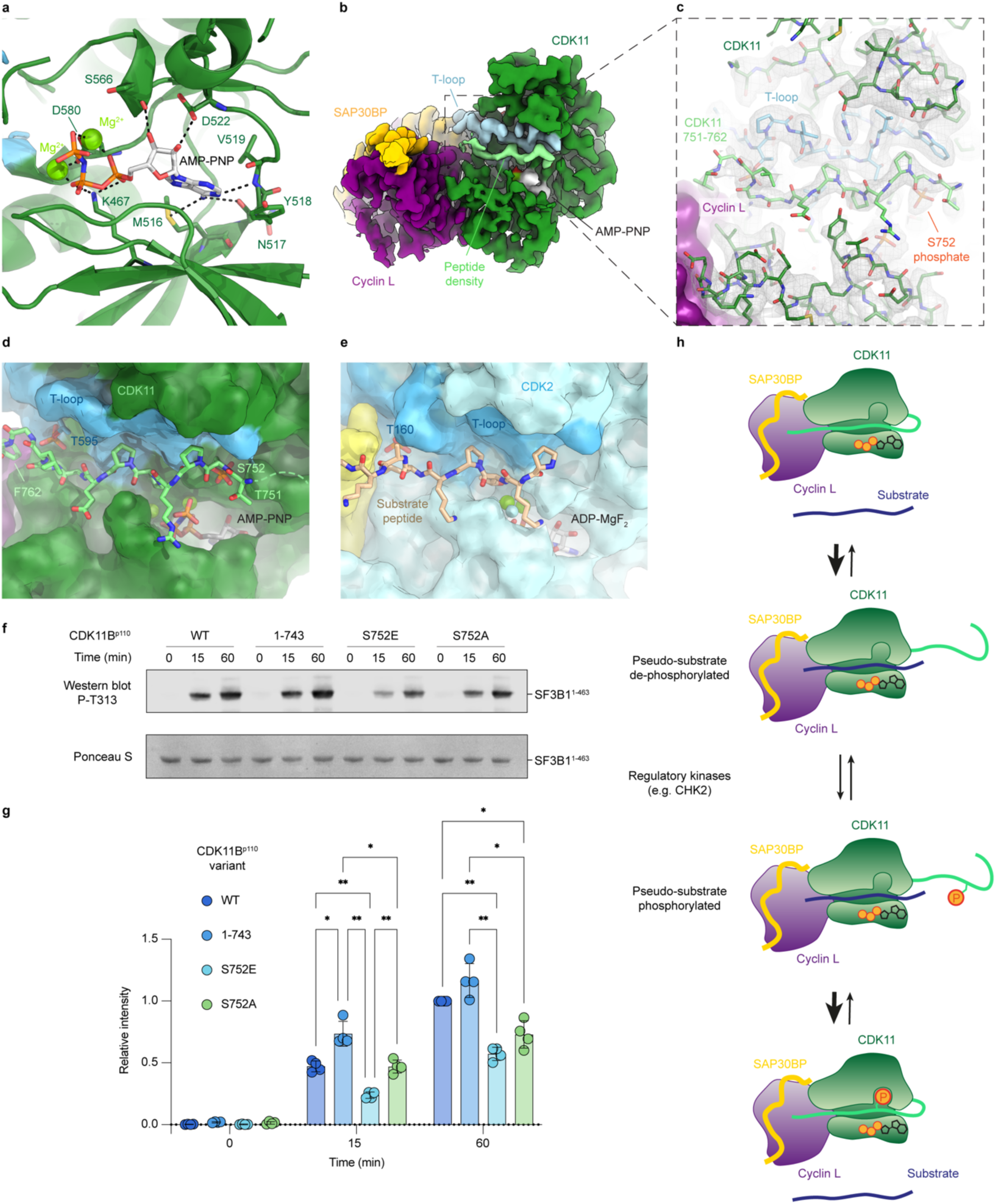
Characterisation of an auto-regulatory pseudo-substrate adjacent to the CDK11 active site. (**a**) Bound AMP-PNP in the active site of CDK11 (green). Residues involved in interactions with the nucleotide are shown as sticks, hydrogen bonds are indicated as black dashed lines, and magnesium ions are shown as green spheres. (**b**) The pseudo-substrate density occupying the substrate binding site of CDK11 is shown in light green (CDK11 green, T-loop blue, SAP30BP yellow, cyclin L purple). (**c**) Cryo-EM map shown as grey mesh and semi-transparent surface, with fitted model coloured as in b. (**d**) Molecular model of the CDK11 pseudo-substrate segment (residues 751-762, light green) occluding the substrate binding site. (**e**) CDK2-cyclin A (cyan and yellow) in complex with a substrate peptide (sand) in the same view as panel C (PDB ID 3QHR) ^34^. (**f**) Kinase assay using an SF3B1^1-463^ substrate and trimeric kinase complexes containing CDK11B^p110^ wild type (WT) and the 1-743, S752E, and S752A variants. SF3B1 phospho-T313 (P-T313) was detected using Western blot. A loading control stained with Ponceau S is provided. (**g**) Quantification of Western blot band intensities, presented as the mean ± standard deviation and individual data points of N = 4 kinase assay technical replicates. The relative activity of the wild-type complex at 60 min was set to 1. The significance level for activity comparisons between complexes, determined by 2-way ANOVA and Tukey’s multiple comparisons test, is provided (*: p < 0.05; **: p < 0.01). (**h**) Schematic illustration of the effect of pseudo-substrate phosphorylation on the efficiency of binding of a simple model substrate to CDK11. Hypothetical models for effects on substrate specificity in the context of more complex substrates are shown in Supplementary Fig. 4e, f. The substrate binding site is schematically represented as a cleft, and the active site is indicated by a stylised ATP molecule.

The cryo-EM map in this region is generally of excellent quality (Supplementary Fig. 2e), allowing unambiguous assignment of side chain positions and molecular contacts. One exception to this is the G-rich loop of CDK11 (residues 445-450, sequence EEGTYG), which exhibits weak density, indicating flexibility. A second element of weak density is positioned just beyond the G-rich loop and the γ-phosphate of the AMP-PNP molecule. While fragmented in fully sharpened cryo-EM maps, this segment becomes continuous in more modestly sharpened maps (sharpening b-factor = -10 Å^2^) and indicates the presence of an extended peptide occupying the substrate binding site of the kinase (Fig. 3b, c). Features of the observed density agree with the identity of CDK11 residues 751-762 (TSPRPPEGGLGF) predicted to occupy this site by AlphaFold3 ^30^ (Fig. 3c). Strikingly, our cryo-EM map and a mass spectrometry analysis of our purified complexes both indicate that S752 is partially phosphorylated in our sample (Fig. 3c, Supplementary Table 1, Supplementary Dataset 1). The phosphate is accommodated in a pocket near the CDK11 active site (Fig. 3c, d). Prior biochemical analysis has shown that the enzyme responsible for phosphorylation of S752 (S737 in isoform 1 studied in ^36^) is checkpoint kinase 2 (CHK2), and that this modification enhances splicing in a cell-based assay ^36^.

The positioning of this CDK11 segment (residues 751-762) coincides with substrate peptides visualised in other CDK structures ^34,37^ and thus precludes substrate access to the active site (Fig. 3d, e). Given that the density for this segment is weaker than for the core fold of CDK11, this structural element is likely dynamic. Overall, these observations indicate that the C-terminal region of CDK11 harbours a pseudo-substrate that might be involved in modulation of substrate access to the active site, suggesting an auto-regulatory role. This is in line with the documented roles of pseudo-substrates in the regulation of kinases ^38^.

To test the effect of this pseudo-substrate on CDK11 activity, we performed kinase assays using wild type CDK11^p110^-cyclin L2-SAP30BP and complexes harbouring the phospho-mimetic CDK11^p110^ S752E mutation, the non-phosphorylatable S752A mutation, and a C-terminal CDK11^p110^ deletion removing the putative pseudo-substrate segment (CDK11B^p110^ ^1-743^; Supplementary Fig. 4a, b). Consistent with a regulatory role of the S752 phosphorylation, the phospho-mimetic S752E mutation reduces kinase activity against the spliceosomal SF3B1^1-463^ substrate (Fig. 3f, g, Supplementary Fig. 4c, d), possibly because the negative charge stabilises the accommodated, active site-blocking, position of the pseudo-substrate. The non-phosphorylatable S752A mutation restores kinase activity to wild type or near-wild type levels (Fig. 3f, g), in agreement with a previous biochemical and cell biological study in which no substantial effect of this mutation on CDK11 kinase activity was found ^36^. Finally, the CDK11B^1-743^ truncation, eliminating the pseudo-substrate, increases CDK11 activity to above wild-type level (Fig. 3f, g). We note that substantial kinase activity is observed in all CDK11 variants tested, indicating that none of the tested configurations of S752 results in complete inhibition, probably because true substrates can outcompete all configurations of the pseudo-substrate, albeit with variable efficiency.

Our data are thus consistent with a regulatory role of this pseudo-substrate within CDK11, which is further modulated by post-translational modification (Fig. 3h). Given our biochemical results showing negative regulation of CDK11 by a phospho-mimetic mutation, it is not immediately clear how S752 phosphorylation would enhance splicing in cellular assays as reported previously ^36^. One hypothesis that might reconcile these observations is that some CDK11 substrates in the splicing pathway might contain binding sites for phosphorylated S752 and the surrounding CDK11 pseudo-substrate sequence. Binding to these substrates would withdraw the pseudo-substrate from its inhibitory position, thereby simultaneously derepressing the kinase and aiding substrate binding (Supplementary Fig. 4e). Alternatively, modulation of the affinity of the pseudo-substrate to the CDK11 substrate binding site could fine-tune which substrates are preferentially phosphorylated by CDK11. S752 phosphorylation could thereby contribute to substrate specificity, in line with proposed models for the role of a phosphorylated pseudo-substrate segment in Casein kinase 1 (ref. ^39^). In the context of this hypothesis, it is important to note that both spliceosomal CDK11 substrates such as SF3B1 (ref. ^16^) and an important transcriptional substrate ^11^, the C-terminal Y_1_S_2_P_3_T_4_S_5_P_6_S_7_ heptapeptide repeat region of the RNA polymerase II subunit RPB1 (the so-called Pol II-CTD) ^40^, contain multiple phosphorylation sites. Pre-existing phosphorylations in these targets may enhance their ability to compete with the phosphorylated pseudo-substrate for CDK11 binding, thereby supporting their further phosphorylation (Supplementary Fig. 4f).

### Cyclins are versatile docking platforms for non-enzymatic CDK modulators

Stably interacting protein factors serve to regulate the assembly and activity of several CDK-cyclin complexes. In addition to the CDK11-cyclin L-SAP30BP complex described here (Fig. 4a-c), these include the metazoan CAK (CDK7-cyclin H-MAT1, Fig. 4d-f) ^41^, the CKM module of mediator (CDK8-cyclin C-MED12-MED13, Fig. 4g-i) ^42^, and pTEF-B (CDK9-cyclin T, Fig. 4j-l), which is captured and effectively hijacked by retroviral proteins, such as HIV Tat ^43^. We performed a structural comparison to determine if shared mechanistic principles or interaction motifs underlie the apparent functional similarities between these accessory factors. Because SAP30BP is primarily a cyclin-binding protein, we did not include exclusively CDK-binding proteins such as Cks1/2 (e.g. ref. ^44^) in this analysis.

**Figure 4.**
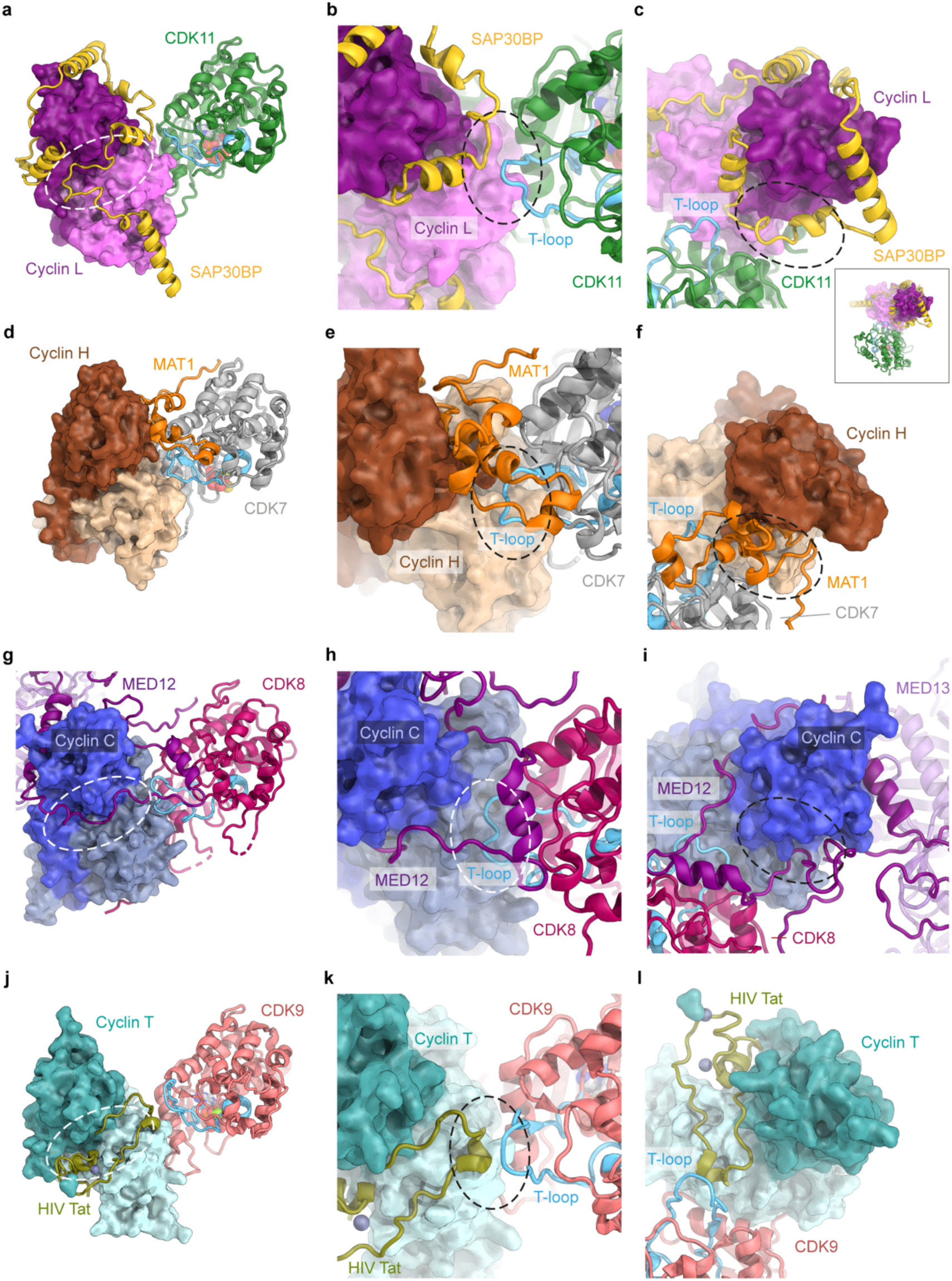
Comparison of CDK-cyclin complexes bound by accessory factors that aid CDK assembly and activity. (**a**-**c**) CDK11-cyclin L-SAP30BP, coloured as in Fig. 2. Throughout the figure, protein subunits are labelled, and areas of interest are denoted by dashed outlines. T-loops are shown in cyan and N- and C-terminal cyclin domains are coloured in distinct shades. An inset shows the complex in the same view as in panel c, but zoomed out for orientation. (**d**-**f**) The human CAK (CDK7-cyclin H-MAT1). CDK7 is shown in grey, MAT1 in orange, and cyclin H in brown. (**g**-**i**) The CKM module of mediator (CDK8-cyclin C-MED12-MED13). CDK8 is shown in red, MED12 in purple, and cyclin C in blue. (**J**-**L**) The HIV protein Tat hijacking the pTEF-B complex (CDK9-cyclin T). CDK9 is shown in salmon, cyclin T in teal, and Tat in olive green.

Our structure of the CDK11-cyclin L-SAP30BP complex reveals that SAP30BP in the CDK11-containing complex, MAT1 in the CDK7-containing CAK complex (PDB ID 6XBZ) ^41^, and MED12 in the CDK8-containing CKM complex (PDB ID 7KPV) ^42^ do not share meaningful structural resemblance even though their functions – complex assembly and enhancing CDK activity – are similar (Fig. 4a, d, g). However, there are some shared interaction regions on their respective CDK and cyclin partners used by SAP30BP, MAT1, and MED12. All three accessory subunits contact the T-loop of their partner CDK (Fig. 4b, e, h) and the region around the first and third conserved helices of the C-terminal cyclin domain of their partner cyclins (Fig. 4c, f, i), albeit in structurally distinct ways. Furthermore, both SAP30BP and MED12 contact the region near the interface between the N- and C-terminal cyclin domains of their partner cyclins (Fig. 4a, g), though again without appreciable similarity in the sequence elements used to form these interactions.

The N-terminal half of the HIV Tat protein also exploits the latter interface region when binding to cyclin T within P-TEFb (PDB ID 3MIA) ^43^, thereby activating CDK9, changing its substrate specificity, and recruiting it to a sequence element within the HIV mRNA to promote transcription ^43^. However, on the molecular level, the details of the interaction again differ from the examples discussed above (Fig. 4j-l). These considerations suggest that CDK-cyclin-binding proteins use a combination of a few shared and additional unique interaction surfaces on their targets, but that the molecular details of these interactions and the CDK-activating interactions employed differ in each case. Our comparison additionally shows that the extent of the interaction interface formed by SAP30BP on cyclin L is uniquely large (3,530 Å^2^), compared to the buried surface area in the other complexes (1,200 Å^2^, 2640 Å^2^, and 1420 Å^2^ for cyclin H-MAT1, MED12-cyclin C, and Tat-cyclin T, respectively). Conversely, the interaction interface between SAP30BP and CDK11 (360 Å^2^) is substantially smaller than for MAT1 and CDK7 (1,220 Å^2^) ^41^. These differences likely reflect the unique function of SAP30BP in stabilisation of cyclin L, whereas MAT1 primarily promotes CDK7-cyclin H complex assembly and activation.

### Structure of CDK11-cyclin L-SAP30BP-OTSG64

OTS964 is a non-covalent, ATP-competitive inhibitor that binds to the active site of its target kinases (Supplementary Fig. 5a). While the structure of the isolated kinase domain of CDK11B in complex with OTS964 has been determined previously (PDB ID 7UKZ) ^29^, results reported in that study indicate that cyclin binding improves OTS964 affinity for CDKs and suggest that the isolated kinase domain might not be able to fully recapitulate the compound-binding properties of the full-length cellular CDK11 protein ^29^. This is in agreement with the general idea that the inhibitor-binding properties of CDKs depend on the state of the regulatory elements of the kinase domain, which are controlled by cyclin binding and T-loop phosphorylation ^45^. Therefore, we decided to investigate the structural basis of the high affinity of OTS964 for CDK11 and its mechanism of selectivity in the context of the activating binding partners of the kinase. To this end, we determined the structure of CDK11-cyclin L-SAP30BP-OTS964 at 2.4 Å resolution (Supplementary Fig. 5a-f).

The pose of OTS964 in our cryo-EM structure of the CDK11-cyclin L-SAP30BP complex (Fig. 5a, Supplementary Fig. 5b) is similar to the previously reported CDK11B-OTS964 X-ray crystal structure (PDB ID 7UKZ) ^29^. The tricyclic ring system of the inhibitor is positioned such as to enable formation of three hydrogen bonds: Two hydrogen bonds to the backbone carbonyl and amide nitrogen of the CDK11 hinge residue V519, and one hydrogen bond either to D522, which is a highly conserved residue among CDKs, or to E445 located within the N-terminal kinase lobe (Fig. 5b). Both D552 and E445 are within hydrogen bonding distance to the exocyclic hydroxy group of OTS964. Considering the expected ionisation state of carboxylic acids at physiological pH, only one of these two possible hydrogen bonds can be formed at a time, and E445 may engage in hydrophobic interactions with the nearby phenyl or dimethylamino groups of the inhibitor instead. D552 being the primary hydrogen bonding partner in this interaction is also consistent with the previous observation that the CDK11 E445G mutation did not impair OTS964 binding ^29^.

**Figure 5.**
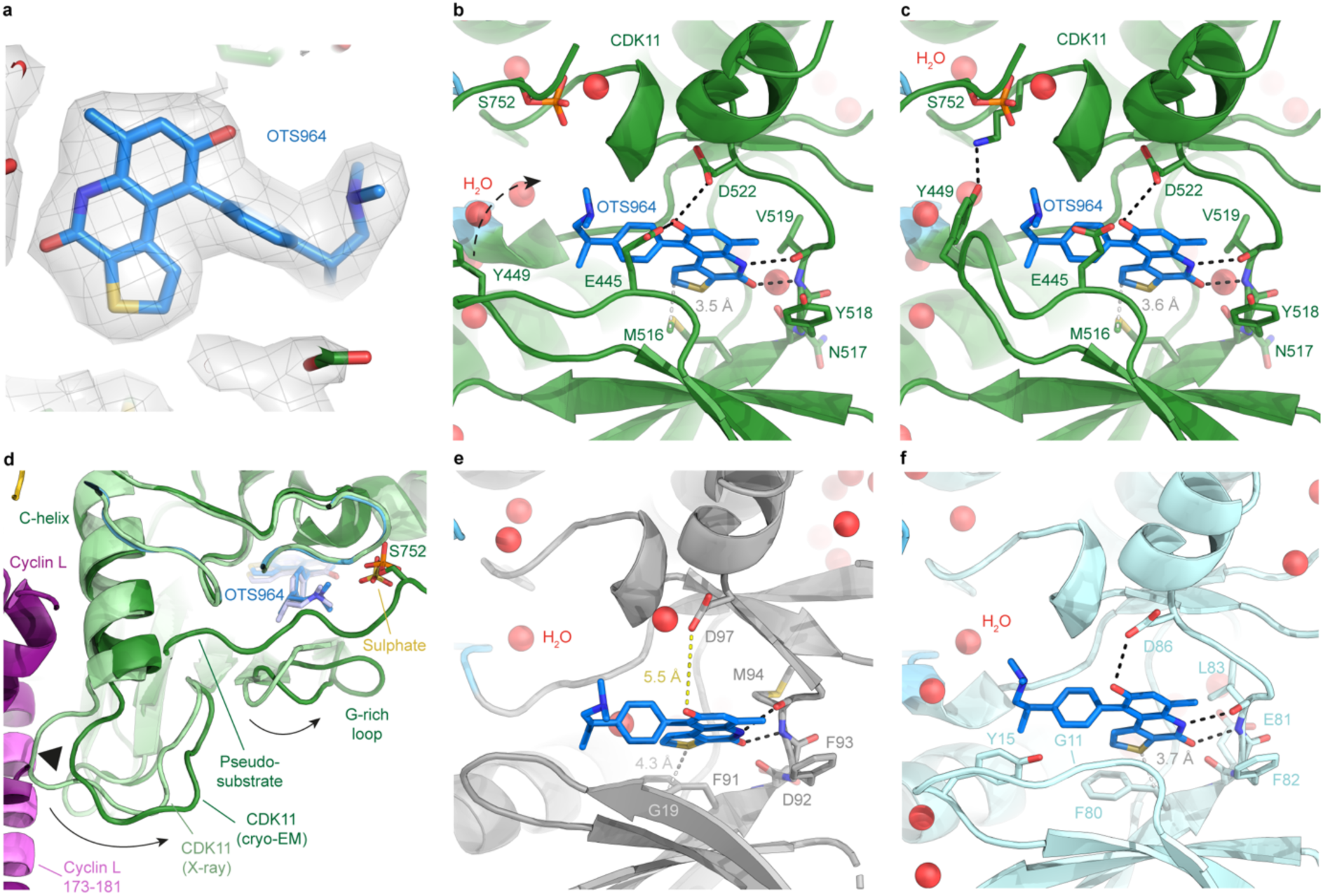
Structure of OTSG64 bound to CDK11-cyclin L-SAP30BP and off-target kinases. (**a**) Rendering of CDK11-bound OTS964 (blue) in the cryo-EM map (shown as semi-transparent grey mesh and surface). (**b**, **c**) Active site views of the CDK11-cyclin L-SAP30BP-OTS964 complex in the two alternative conformations observed in the cryo-EM reconstruction. Hydrogen bonds (donor-acceptor distance 3.5 Å or less) are indicated by black dashed lines. The shortest distance between inhibitor and gatekeeper residue (M516 in CDK11) is indicated and shown by a white dashed line. The change in position of Y449 between panels b and c is indicated with a dashed arrow in panel b. (**d**) Comparison of the structure of CDK11-cyclin L-SAP30BP-OTS964 (CDK11 in green, cyclin L in purple with residues 173-181 in pink, OTS964 in blue) to the structure of isolated CDK11B-OTS964 (PDB ID 7UKZ ^29^; light green and light blue, respectively). The site of steric incompatibility of the isolated CDK11B conformation with the presence of cyclin L is indicated by a black triangle. Shifts in adjacent loops because of cyclin binding are indicated by arrows. (**e**) Active site view of the CAK-OTS964 complex (CDK7 is shown in grey, water molecules as red spheres). Only the hydrogen bonds to the hinge region are preserved because several hydrogen bonding partners are absent or beyond hydrogen bonding distance (distance to D97 indicated by a yellow dashed line). The distance to the gatekeeper residue (F91 in CDK7) is enlarged (yellow dashed line). (**f**) Active site view of the CDK2-cyclin A2-OTS964 complex (CDK2 is shown in cyan). OTS964 forms three hydrogen bonds to CDK2, as it is within hydrogen bonding distance of CDK2 D86. Y15 is tucked under the dimethylamino-propyl group of OTS964.

The density for the G-rich loop in the OTS964-bound complex is weaker than neighbouring regions near the CDK11 active site. Interestingly, our cryo-EM density suggest that two alternative conformations of this loop exist. One of the conformations brings CDK11 residue Y449 to within hydrogen bonding distance of CDK11 K564, thereby bookending the dimethylamino group of OTS964 (Fig. 5c, Supplementary Fig. 5g, h). It is worth noting that phosphorylation of residues equivalent to Y449 in other CDKs (Y15 in CDK1/2) are known to serve to inactivate these kinases ^2^. However, our mass spectrometry analysis (Supplementary Table 1, Supplementary Dataset S1) did not detect phosphorylation of Y449 or the neighbouring residue T448 (T14 in CDK1/2). This suggests that the alternative G-loop conformations observed in our structure may be independent of post-translational modifications of this region.

Compared to the X-ray crystal structure of the isolated CDK11B-OTS964 complex, there are modest motions within the CDK11 kinase domain upon association with cyclin L and SAP30BP (Fig. 5d). The motions of the αC-helix are minimal, but due to steric interference with cyclin L residues 173-181, the loop preceding the αC-helix is pushed towards the active site, which in turns impacts the conformation of the G-rich loop, with possible implications for binding of inhibitors and nucleotides. Importantly, the G-rich loop conformation in which Y449 points towards the inhibitor and K564 (Fig. 5c) was not observed in the X-ray crystal structure. Therefore, formation of the CDK11-cyclin L complex may influence inhibitor binding by affecting the conformational space that loop regions in the N-terminal kinase domain can access.

The pseudo-substrate segment of CDK11 is still visualised in the inhibitor-bound structure (Fig. 5b-d, Supplementary Fig. 5i), ruling out that nucleotide binding is required for it to assume this position. Interestingly, the position of the phosphate moiety in phosphorylated S572 coincides with the position of a sulphate molecule visualised in the CDK11-OTS964 X-ray crystal structure (PDB ID 7UKZ) ^29^, further supporting the assignment of this density to a similarly negatively charged phosphate in our structure (Fig. 5d).

### Mechanism of OTSG64 selectivity for CDK11 over other CDKs

Due to the high sequence and structural homology among the CDK family, discovery of inhibitors that specifically target a single CDK poses a substantial challenge ^1,35^. Therefore, we determined the structures of CAK-OTS964 and CDK2-cyclin A-OTS964, representing possible off-target complexes, to obtain further insight into OTS964 selectivity for CDK11 relative to other CDKs (Fig. 5e, f, Supplementary Fig. 6a-n). The quality of the density of the inhibitor in the CDK7-containing CAK-OTS964 complex cryo-EM map at 2.3 Å overall resolution is lower compared to that in the CDK11-containing target complex. This is particularly pronounced for the dimethylamino-propan-2-yl-phenyl substituent extending from the aromatic core of the compound (Supplementary Fig. 6k, l). This suggests that the inhibitor might exhibit conformational flexibility within the CDK7 active site, rather than being stably bound with high affinity in a single pose as observed in the CDK11 complex. This is in line with the poor ability of OTS964 to inhibit CDK7 (Supplementary Fig. 7a, b). The active sites of CDK7 and CDK11 differ in some of the residues that form contacts to OTS964 in the CDK11 complex, providing a possible explanation for this observation. Most notably, CDK11 residue E445 is equivalent to G19 in CDK7, which leads to loss of any inhibitor interactions involving this residue in CDK7. Additionally, CDK7 D97 and F91 are further away from the inhibitor than the equivalent residues M516 and D522 in the CDK11 complex, which may further destabilise binding of OTS964 due to lack of hydrophobic packing and hydrogen bonding interactions, respectively (Fig. 5e). Overall, the poor affinity of OTS964 for CDK7 is explained by the absence of several key interactions due to amino acid differences between CDK7 and CDK11 and lack of shape complementarity of the CDK7 active site.

The differences in the inhibitor interactions between the CDK2-cyclin A-OTS964 complex, determined at 2.5 Å resolution, and the CDK11-cyclin L-SAP30BP-OTS964 complex are considerably smaller than those observed for the CDK7-bound inhibitor complex. The cryo-EM density for CDK2-bound OTS964 is of similar quality as for the CDK11-bound complex (Supplementary Fig. 6m, n). In the context of CDK2, OTS964 can form hydrogen bonds to the hinge region as well as to CDK2 D86 (equivalent to CDK11 D522, Fig. 5f). CDK2 Y15 is tucked under the dimethylamino-propyl group of OTS964. This conformation has been observed in inhibitor-bound CDK2 complexes before ^46^, but it is incompatible with nucleotide binding ^47^. In this position, Y15 might contribute to inhibitor binding by allowing burial of additional surface area and formation of van-der-Waals contacts. One difference between the CDK2 and CDK11 active sites is that CDK2 carries a glycine in the position of CDK11 E445. The CDK2 G11E mutation has been observed to increase OTS964 affinity for CDK2 ^29^. Our structure suggests that the glutamate residue introduced in this CDK2 mutant might be able to form either a hydrogen bond to the exocyclic hydroxy group of OTS964, an electrostatic interaction with the protonated dimethylamino-propyl group of the inhibitor, or hydrophobic contacts with the phenyl and dimethylamino groups of the inhibitor. All these possibilities would explain increased OTS964 binding to the CDK2 G11E mutant. Given that the hydrogen bonding ability of the exocyclic hydroxy group of OTS964 is likely satisfied by interactions with CDK2 D86, hydrophobic packing to nearby portions of the inhibitor is the most likely of these mechanisms ^29^.

Further contributions to selectivity may arise from shape complementarity, considering that the inhibitor moves slightly towards the gatekeeper (the slightly more recessed F80 in CDK2 instead of M561 in CDK11) in the CDK2-bound structure (Supplementary Fig. 7c). Finally, the identity of the residue in the position of CDK11 G579, a position in which CDK2 harbours an alanine (A144), is known to affect inhibitor binding. Prior work has shown that the introduction of larger, sterically incompatible residues, such as serine in CDK11 G579S, leads to strong resistance to OTS964 (refs ^16,28,29^). Our structures show that the backbone conformation at this position differs between OTS964-bound CDK11 and CDK2 (Supplementary Fig. 7c). Due to this conformational difference, CDK2 A144 does not protrude further towards the inhibitor than CDK11 G579 does (Supplementary Fig. 7c). Unexpectedly, CDK11 G579 and CDK2 A144 in the nucleotide-bound state ^47^ assume overlapping backbone conformations because of a conformational switch in CDK11 G579 (Supplementary Fig. 7d). It is thus possible that the specific ability of CDK11 G579 to undergo this conformational change in the inhibitor-bound state leads to improved packing or shape complementarity, thereby supporting OTS964 binding. This hypothesis would account for the partial OTS964 resistance of cells harbouring the CDK11 G579A mutation ^28,29^, which should otherwise be accommodated without steric clashes due to the small size of the alanine side chain ^29^. However, prior data showing that the G579A mutation enhances, rather than reduces, OTS964 binding to isolated CDK11 *in vitro* ^29^ indicate that the cellular environment or the assembly of CDK11 into cyclin-bound complexes may have additional, more subtle, effects on inhibitor binding.

### Conclusion

Taken together, our structural and biochemical data provide detailed insight into the molecular mechanisms governing CDK11 activity, regulation, and inhibition. Our finding that cyclin L2 on its own is unstable – in agreement with prior results ^17,29^ – but becomes stabilised when expressed with its SAP30BP partner indicates that SAP30BP is a general cofactor for CDK11-cyclin L, in addition to its more specific functions in certain splicing pathways (Fig. 6a). This explains the unusually extensive but highly specific interaction interface between cyclin L and SAP30BP. Furthermore, we have characterised a pseudo-substrate within the C-terminal region of CDK11, indicating that substrate access is modulated by post-translational modifications of this segment (Fig. 3h). This has possible implications for regulation of substrate specificity, though the specific mechanisms by which CDK11 pseudo-substrate modifications affect its biological function remain to be established (Supplementary Fig. 4e, f). We note that phosphorylation of S752 within this pseudo-substrate segment has previously been linked to CDK11 dimerization rather than regulation of kinase activity ^36^. While S752 phosphorylation might be able to promote the reciprocal binding of two CDK11 molecules to each other’s C-terminal segments, analogous to a domain-swap (Fig. 6b), our structural and biochemical results are more consistent with a pseudo-substrate function of this sequence (Fig. 6c). Finally, our structures of inhibitor-bound kinase complexes provide insight into OTS964 specificity for CDK11 and uncover conformational differences between the fully assembled CDK11-cyclin L-SAP30BP complex and the isolated CDK11B kinase domain. Thereby, we provide a structural framework for rational design and discovery of next-generation cancer therapeutics targeting CDK11.

**Figure 6.**
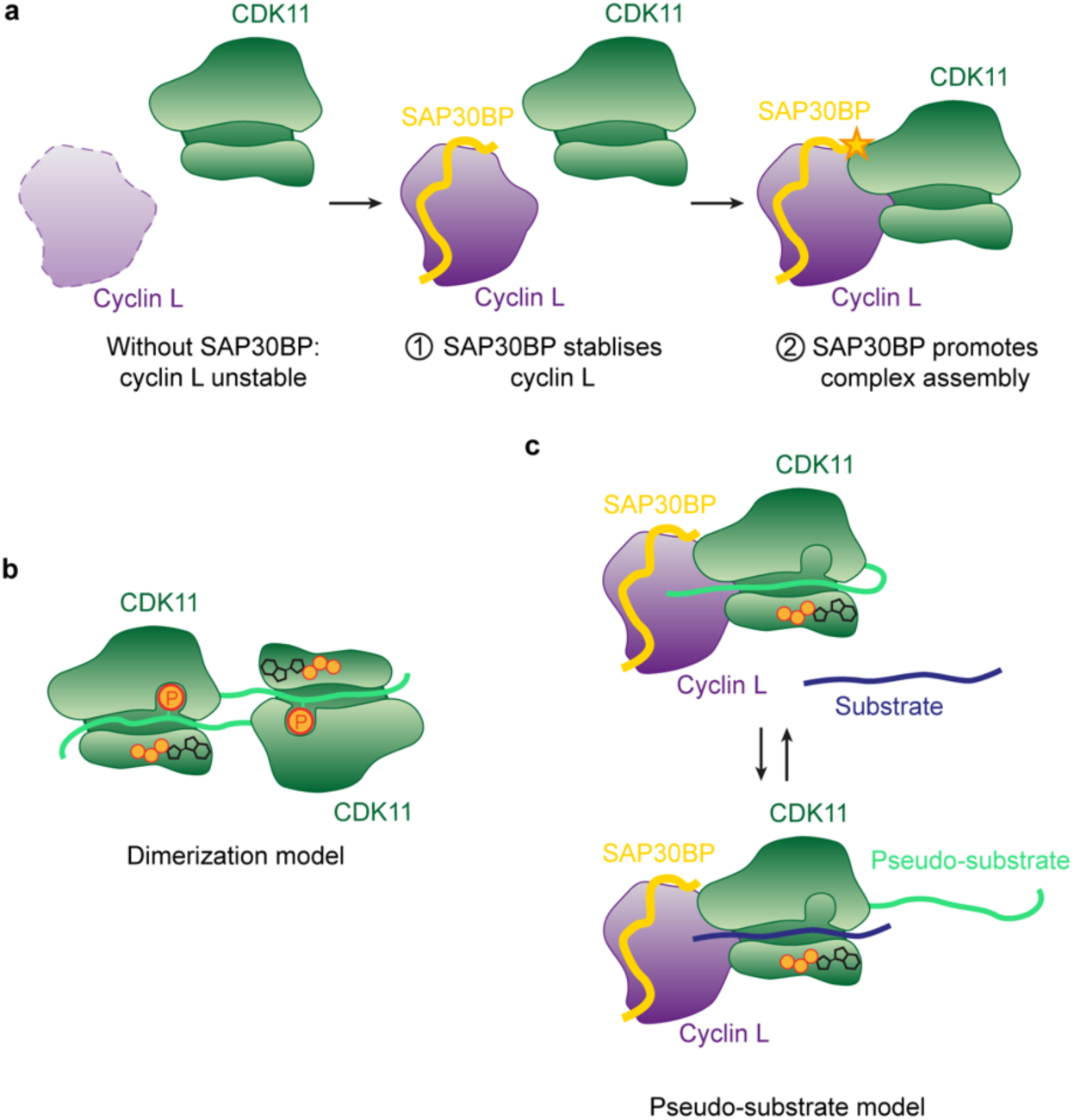
Roles of SAP30BP and the CDK11 pseudo-substrate in complex assembly and kinase regulation. (**a**) Schematic illustration of the dual role of SAP30BP in CDK11 activation by (1) stabilising cyclin L and (2) enhancing complex formation with CDK11. CDK11 and cyclin L are represented by green and purple shapes, and SAP30BP is represented by a yellow ribbon. (**b**) Model of how pseudo-substrate phosphorylation might lead to CDK11 dimerization. (**c**) Schematic illustration of the pseudo-substrate model applied to the binding of a simple model substrate to CDK11. The substrate binding site is schematically represented as a cleft, and the active site is indicated by a stylised ATP molecule.

## METHODS

### Protein expression and purification

#### CDK11-cyclin L-SAP30BP

Synthetic, codon optimised genes encoding CDK11B, cyclin L2, and SAP30BP (Twist Bioscience) were cloned into 438-series vectors for baculovirus-based expression in insect cells ^48^, resulting in a multi-protein construct expressing Twin-Strep-tagged CDK11B (357-795), His_6_-tagged Cyclin L2 (1-319), and MBP-tagged SAP30BP (Supplementary Fig. 1a, b). Assembled constructs were transformed into EMBacY cells for transposition into baculoviral bacmids ^49^. Bacmids were prepared by isopropanol precipitation before transfection into Sf9 (*Spodoptera frugiperda*) insect cells (Thermo Fisher Scientific) using Insect GeneJuice Transfection Reagent (Sigma-Aldrich). After two rounds of baculovirus amplification, 20 mL of insect cell culture supernatant was used to infect 1-L cultures of High5 (*Trichoplusia* ni) insect cells (Thermo Fisher Scientific). After 72 h of incubation, insect cells were harvested and stored at -80 °C for later use.

The CDK11-cyclin L-SAP30BP complex was purified by Strep-Tactin-affinity and gel filtration chromatography. Cells were thawed and resuspended in purification buffer (250 mM KCl, 2 mM MgCl_2_, 40 mM HEPES-KOH pH 7.9, 5 mM β-mercaptoethanol, 10% (v/v) glycerol) supplemented with protease inhibitors and DNase I before lysis by successive cycles of freeze-thaw. Cell debris were pelleted by centrifugation at 18,000 rpm for 30 min at 4°C in a JA-25.50 rotor (Beckman Coulter), and cleared lysate was incubated for 45 min with 2.5 mL of Strep-Tactin Superflow Plus resin (Qiagen). Beads were washed in purification buffer and purified protein was eluted in purification buffer supplemented with 10 mM desthiobiotin (Sigma-Aldrich). Affinity tags were cleaved by Tobacco Etch Virus (TEV) protease at 4°C for 1 h before proceeding directly to size exclusion chromatography, performed on a Superdex 200 Increase 10/300 GL column (GE Healthcare). Peak fractions were pooled, concentrated to approx. 2 mg/mL, snap frozen in liquid N_2_, and transferred to -80°C for long-term storage.

#### CDK11 and cyclin L-SAP30BP for pull-down assays

An expression construct encoding Twin-Strep-tagged CDK11B (residues 357-795) was cloned and used to prepare bacmids, which were then transfected into Sf9 insect cells (Thermo Fisher Scientific). Two multi-expression constructs were also cloned for expression of His_6_-tagged Cyclin L2 (residues 1-319) and either the WT or 4A mutant (K157A, R160A, N161A, K167A) of MBP-tagged SAP30BP, and similarly used to prepare bacmids. After transfection and two rounds of baculovirus amplification, 20 mL (Cyclin L2-SAP30BP) or 40mL (CDK11B) of insect cell culture supernatant was used to infect 1-L (cyclin L2-SAP30BP) or 2-L (CDK11B) cultures of High5 insect cells (Thermo Fisher Scientific). After 72 h of incubation, insect cells were harvested and stored at -80 °C for later use.

Strep-tagged CDK11B (357-795) was purified as described for the CDK11-cyclin L-SAP30BP complex above, only omitting the cleavage of the affinity tags. WT and mutant cyclin L2 (1-319)-SAP30BP complexes were purified by Nickel-affinity and gel filtration chromatography. Cells were thawed and resuspended in purification buffer (250 mM KCl, 2 mM MgCl_2_, 40 mM HEPES-KOH pH 7.9, 5 mM β-mercaptoethanol, 10% (v/v) glycerol, 10 mM imidazole) supplemented with protease inhibitors and DNase I before lysis by successive cycles of freeze-thaw. Cell debris were pelleted by centrifugation at 18,000 rpm for 30 min at 4°C in a JA-25.50 rotor (Beckman Coulter), and cleared lysate was incubated for 45 min with 2.5 mL of Ni-NTA Superflow resin (Qiagen). Beads were washed in wash buffer (purification buffer with 25 mM imidazole) and purified protein was eluted in elution buffer (purification buffer with 300 mM imidazole). Affinity tags were cleaved by TEV protease at 4°C for 1 h before proceeding directly to size exclusion chromatography, performed on a Superdex 200 Increase 10/300 GL column (GE Healthcare). Peak fractions were pooled, concentrated to approx. 2 mg/mL, snap frozen in liquid N_2_, and transferred to -80°C for long-term storage.

#### CDK11-cyclin L-SAP30BP complexes for kinase assays

CDK11^p110^ complexes for kinase assays were expressed in insect cells. Point mutations were cloned into an existing construct of CDK11B^p110^ in a 438-series vector, and the truncation mutant (1-743) was generated through introduction of a premature stop codon. Complete complexes were obtained by infecting Sf9 insect cells with bacmid DNA to generate separate baculoviruses encoding either Strep-tagged-CDK11B^p110^ (WT or mutant) or His_6_-tagged-cyclin L2 and MBP-tagged-SAP30BP. After two rounds of baculovirus amplification, these were combined for co-expression in High5 insect cells by inoculating two 0.5-L cultures with 10 mL of each baculovirus. After 72 h of incubation at 27 °C, cultures were harvested and pellets stored at -80 °C for later use.

WT and mutant complexes were purified by Strep-Tactin affinity and gel filtration chromatography as described for the CDK11-cyclin L-SAP30BP complex used for structure determination, with affinity tags cleaved by incubation with TEV protease. Peak fractions were pooled, concentrated to approx. 1 mg/mL, snap frozen and stored at -80 °C for later use.

#### SF3B1 for kinase assays

GST-SF3B1 (SF3B1 residues 1-463) was expressed from a synthetic, codon-optimised plasmid in a pGEX-4T-1 backbone (Genscript) in bacterial *E. coli* BL21-RIL cells. Two 10 mL cultures of LB containing 100 µg/mL ampicillin and 34 µg/mL chloramphenicol were inoculated with a single colony of BL21-RIL cells transformed with pGEX-4T-1-SF3B1^1-463^ and grown overnight at 37 °C with vigorous shaking. Two 1-L cultures of LB containing 100 µg/mL ampicillin and 34 µg/mL chloramphenicol were each inoculated with the entirety of a 10 mL overnight culture and incubated at 37 °C until the optical density (OD) had reached 0.6. Cultures were induced with 0.5 mM isopropyl-thiogalactoside (IPTG) and incubated at 37°C for a further 4 hours. Cultures were harvested by spinning at 6000 x g for 18 min at 4°C using a JLA 8.1 rotor and pellets were snap frozen in liquid nitrogen and stored at -80°C for later use.

GST-SF3B1^1-463^ was purified by GST-affinity resin. Pellets were thawed and resuspended in purification buffer (150 mM NaCl, 2 mM MgCl_2_, 50 mM HEPES-KOH pH 7.5, 5 mM β-mercaptoethanol, 10% (v/v) glycerol) supplemented with protease inhibitors and DNase I before lysis by sonication. Cell debris were pelleted by centrifugation at 18,000 rpm for 30 min at 4°C in a JA-25.50 rotor (Beckman Coulter), and cleared lysate was incubated for 45 min with 2.5 mL of Glutathione Sepharose 4B resin (Cytiva) at 4 °C. Beads were washed in purification buffer and purified protein was eluted in purification buffer supplemented with 10 mM reduced glutathione (Sigma). Eluted fractions were spun at 20,000 x g to pellet aggregates, aliquoted, snap frozen and stored at -80 °C.

#### CDK7-cyclin H-MAT1(Δ21G)

Structure determination of the CAK-OTS964 complex employed a construct lacking the N-terminal, flexibly attached domain of MAT1 ^41^, removal of which had previously been observed to improve the orientation distribution of CAK particles in cryo-EM experiments ^50^. The complex was expressed and purified as described above for CDK11-cyclin L-SAP30BP, except that a nickel affinity purification step using Ni-NTA Superflow beads (Qiagen) was added before incubation with the Strep-Tactin Superflow resin, and that the cleaved tags and TEV protease were removed by incubation with Ni-NTA Superflow beads before gel filtration.

#### CDK2-cyclin A2

CDK2 and cyclin A2 were expressed separately in Rosetta DE3 pLysS cells. CDK2 was expressed with an N-terminal His_10_-tag (Addgene plasmid #79726; http://n2t.net/addgene:79726; RRID:Addgene_79726, provided by John Chodera, Nicholas Levinson, and Markus Seeliger) ^51^, and cyclin A2 was expressed with an N-terminal His_6_-tag. The cyclin A2 sequence was provided by Jonathon Pines and the 15-H bacterial expression vector was a gift from Avinash Patel. For protein expression, Rosetta DE3 pLysS cells harbouring the individual expression constructs were grown at 37°C to OD ∼0.6, induced with 1 mM IPTG, and then incubated at 24°C overnight. The cultures were harvested by centrifugation, frozen in liquid nitrogen, and stored at -80°C.

For protein purification, cell pellets of CDK2- and cyclin A2-expressing cells were combined and resuspended in purification buffer (50 mM HEPES pH 7.5, 180 mM NaCl, 5 % (v/v) glycerol, 2 mM MgCl_2_, 2 mM DTT) supplemented with protease inhibitors and DNase I. The cells were lysed by sonication and the lysate was clarified by two centrifugation steps at 18,000 rpm in a JA-25.50 rotor at 4°C. The cleared lysate was supplemented with 10 mM imidazole and loaded onto a 5 mL HisTrap column (Cytiva), which was then attached to an Akta Pure M25 instrument (Cytiva), washed using purification buffer supplemented with 20 mM imidazole, and eluted using a linear imidazole gradient from 20-300 mM across 12 column volumes. The column was further eluted using 30 mL of buffer at 300 mM, and fractions containing both proteins were combined, concentrated to 4 mg/mL, and incubated with TEV protease at 4°C overnight to cleave off the His_6/10_ tags. After incubation with Ni-NTA Superflow beads (Qiagen) for removal of the cleaved tags and the TEV protease, the supernatant was concentrated to 1.5 mL, frozen in liquid nitrogen, and stored overnight. This frozen protein aliquot was thawed, and the protein complex was then further purified using a Superdex 200 Increase 10/300 column (Cytiva). Peak fractions were combined, yielding a concentration of approx. 2 mg/mL, and flash frozen in liquid nitrogen for storage at -80°C.

### Pull-down assay with and without cyclin L-SAP30BP co-expression

Cloning was performed to generate 438-series vectors for insect cell expression encoding His_6_-tagged cyclin L2, MBP-tagged SAP30BP and Strep-tagged CDK11B^p110^. A multi-protein co-expression construct was also made containing all three subunits with their respective affinity tags in one vector. These constructs were used to prepare bacmids and then transfected into Sf9 insect cells to generate baculoviruses. After one round of baculovirus amplification, a 50 mL culture of Sf9 cells was inoculated for each co-expression test. For expressions of cyclin L2 alone, or cyclin L2 with one other protein subunit, the culture was inoculated with 1 mL of each virus encoding a protein (1-2 mL). For the co-expression of all three proteins, 1 mL of baculovirus encoding the single multi-protein construct was added to the culture. Inoculated cultures were incubated at 27 °C for 72 hours and cell pellets were harvested by spinning at 1,000 x g for 15 min, before being stored at -80 °C for later use.

The pull-down assay was performed using the nickel-binding capability of His_6_-tagged cyclin L2. Pellets were resuspended in 3 mL of assay buffer (250 mM KCl, 2 mM MgCl_2_, 40 mM HEPES-KOH pH 7.9, 5 mM β-mercaptoethanol, 10% (v/v) glycerol, 10 mM imidazole) and 1.25 mL of this resuspension was then used in the pull-down. Resuspended cells were lysed by three successive freeze-thaw cycles and the lysate cleared by spinning at 20,000 x g for 10 min at 4 °C. Cleared lysate was incubated with 80 µL Ni-NTA Superflow resin for 15 min at 4 °C. Three washes were performed with 1 mL of assay buffer (supplemented with imidazole to 25 mM). Bound proteins were eluted in 100 µL of assay buffer (supplemented with imidazole to 300 mM).

### Pull-down assay with cyclins H, K, and T

Synthetic, codon-optimised genes for cyclins H, K and T were cloned into 438-series vectors for insect cell expression, resulting in constructs encoding each cyclin with an N-terminal His_6_ tag. An additional construct was prepared encoding MBP-tagged SAP30BP. Constructs were transformed into EMBacY cells for transposition into baculoviral bacmids, and the resulting bacmids were used to transfect Sf9 cells for production of baculovirus. After one round of baculovirus amplification, pull-down co-expressions were made by infecting 50 mL cultures of Sf9 insect cells with 1 mL each of virus encoding MBP-tagged SAP30BP and one of the His_6_-tagged cyclin variants. After cultures were incubated for 72 h at 27 °C, pellets were harvested by centrifugation at 1000 x g for 10 min and stored at -80 °C for later use.

Separate pull-down assays were performed using Nickel and Amylose affinity resin. Co-expression cell pellets were resuspended in 3 mL assay buffer (250 mM KCl, 2 mM MgCl_2_, 40 mM HEPES-KOH pH 7.9, 5 mM β-mercaptoethanol, 10% (v/v) glycerol) and this was then divided into 1.5 mL for each of the different affinity pull-downs. The Nickel pull-down suspension was supplemented with imidazole to a concentration of 10 mM. Cells were lysed by 3 successive freeze-thaw cycles of resuspended pellets, and lysate was cleared by spinning at 20,000 x g for 15 min at 4 °C. Cleared lysate was incubated with 200 µL Ni-NTA Superflow resin or 500 µL Amylose resin, rotating for 30 min at 4 °C. Three 1 mL washes were performed using assay buffer (supplemented with 25 mM imidazole for Ni affinity pull-downs). Bound proteins were eluted in 100 µL of assay buffer supplemented with 300 mM imidazole (nickel-affinity pull-downs) or 200 µL of assay buffer supplemented with 10 mM maltose.

### Kinase assays for pseudo-substrate characterisation

50 nM or 66 nM CDK11^p110^-cyclin L-SAP30BP complex (WT or mutant) was incubated with 6 µM GST-SF3B1^1-463^ and 2 mM ATP in a total volume of 100 µL, in assay buffer (20 mM HEPES-KOH pH 7.9, 5 mM MgCl_2_, 2 mM DTT, 2.5% (v/v) glycerol) for 60 min at RT. Samples were taken at 0-, 15- and 60 min time points, diluted 1:10 in assay buffer, and quenched by combination with SDS-loading dye. N = 4 technical replicates, each measured from a distinct kinase reaction, were performed for each CDK11 variant complex. 15 µL samples were analysed by SDS-PAGE using Nu-PAGE 4-12% Bis-Tris gels such that the 0-, 15- and 60-min samples from all four complexes were run on a single gel, for each repeat.

Western blotting was performed following transfer to nitrocellulose membranes. Membranes were stained using Ponceau S and imaged using the ChemiDoc imager to obtain loading controls. Membranes were destained in PBST (1x phosphate buffered saline (PBS) supplemented with 0.1% v/v Tween 20), then blocked in 5% BSA/PBST (5% w/v bovine serum albumin dissolved in PBST) at 4 °C overnight. Primary incubation was performed for 1 h at RT using a rabbit anti-phospho-SF3B1 (Thr313) primary antibody (Cell Signalling Technology, cat. #25009) diluted 1:2000 in 5% BSA/PBST. Membranes were washed in PBST before incubation with a goat anti-rabbit secondary antibody conjugated to IRDye 800CW (LICORbio, cat. 926-32211) diluted 1:20,000 in 5% BSA/PBST. Membranes were washed in PBST, then imaged using the Odyssey CLx imaging system (LICORbio), using the 800 nm channel.

Band intensities were quantified using ImageJ ^52^ and blot bands were normalised to those of loading controls (SF3B1 substrate). The WT signal was set to a relative intensity of 1 for each replicate. Statistical analysis of N = 4 technical replicates was performed by 2-way ANOVA and Tukey’s multiple comparisons test, as implemented in PRISM (version 10.4.1; GraphPad Software).

### Enzyme inhibition assays

*In-vitro* kinase assays of OTS964 with CDK11-cyclin K, CDK2-cyclin A2, and CDK7-cyclin H-MAT1 were performed by Reaction Biology GmbH (Freiburg, Germany). ATP concentrations were set to the apparent K_m_ of the respective enzyme. Raw data were normalized using DMSO (negative) and enzyme-free (positive) controls on the same plates. Data were analysed in PRISM (version 10.3; GraphPad Software) as inhibitor concentration vs. normalized response with variable slope. Asymmetric confidence intervals were computed at a significance level of 95%.

### Phosphoproteomics analysis

The CDK11^p110^-SAP30BP-cyclin L2 sample (approx. 24 μg) was adjusted with triethylammonium bicarbonate buffer (TEAB) at a final concentration of 100 mM. Proteins were reduced and alkylated with 5 mM tris-2-carboxyethyl phosphine (TCEP) and 10 mM iodoacetamide (IAA) simultaneously for 60 min in the dark and digested overnight with trypsin at a final concentration of 50 ng/μL (Pierce). The sample was dried and peptides were cleaned up with desalting spin columns (Pierce) followed by phosphopeptide enrichment with the High Select Fe-NTA kit (Pierce) according to manufacturer’s instructions. Both the eluate and flow-through solutions were kept for LC-MS analysis.

LC-MS analysis was performed on a Dionex UltiMate 3000 UHPLC system coupled with an Orbitrap Ascend Mass Spectrometer (Thermo). Peptides were reconstituted in 30 μL 0.1% TFA and 15 μL (eluate) or 3 μL (flow-through) were loaded to the Acclaim PepMap 100, 100 μm × 2 cm C18, 5 μm trapping column at 10 μL/min flow rate of 0.1% TFA loading buffer. Peptides were then subjected to a gradient elution on a capillary column (Waters, nanoE MZ PST BEH130 C18, 1.7 μm, 75 μm × 250 mm) connected to the EASY-Spray source at 45 °C with an EASY-Spray emitter (Thermo, ES991). Mobile phase A was 0.1% formic acid and mobile phase B was 80% acetonitrile, 0.1% formic acid. The separation method at a flow rate of 300 nL/min was an 80 min gradient from 3%-26% B. Precursors between 375-1,500 m/z and charge states 2-7 were selected at 120,000 resolution in the top speed mode in 3 sec and isolated for HCD fragmentation (collision energy 32%) with quadrupole isolation width 0.7 Th, Orbitrap detection with 30,000 resolution and 59 ms Maximum Injection Time. Targeted MS precursors were dynamically excluded from further isolation and activation for 45 seconds with 10 ppm mass tolerance.

Peptide and protein identification was conducted in Proteome Discoverer 3.0 (Thermo Fisher Scientific) with the Sequest HT search engine. Precursor and fragment mass tolerances were 20 ppm and 0.02 Da respectively with a maximum of 2 trypsin missed cleavages allowed. Dynamic modifications included Carbamidomethyl at C, Oxidation of M, Deamidation of N/Q, and Phosphorylation of S/T/Y. Spectra were searched against a FASTA file containing reviewed *Homo sapiens* UniProt entries. Peptides were filtered at q-value < 0.01 using the Percolator node and target-decoy database search. Phosphorylation localization probabilities were estimated with the IMP-ptmRS node.

### Cryo-EM sample preparation

Cryo-EM samples containing CDK11-cyclin L-SAP30BP were prepared on UltrAuFoil R1.2/1.3 holey gold grids (Quantifoil Microtools). Purified protein complex was diluted to approx. 0.4 mg/mL in dilution buffer (200 mM KCl, 2 mM MgCl_2_, 40 mM HEPES-KOH pH 7.9) and combined with 1 mM AMP-PNP (Sigma-Aldrich) or 25 µM OTS964 (MedChemExpress; dissolved at 25 mM in 100% DMSO), in the presence of a 200 µM of a 21-mer cryo-peptide (sequence MALKVTKNSKINAENKAKINM) on ice. 4 µL of sample were applied to plasma cleaned grids (Tergeo-EM plasma cleaner, PIE Scientific), mounted in a Vitrobot Mark IV (Thermo Fisher Scientific) operated at 5°C and 100 % humidity, blotted for 1, 1.5 or 2 sec, and plunged into liquid ethane at liquid N_2_ temperature. Subsequently, the grids were clipped into autogrid cartridges (Thermo Fisher Scientific) compatible with the Glacios and Krios autoloader systems.

Cryo-EM samples of CAK-OTS964 and CDK2-cyclin A-OTS964 complexes were prepared as described above, except that 50 µM OTS964 were used to overcome the lower affinity for the off-target kinases. No peptide additive was used for the CAK-containing complex.

### Initial cryo-EM screening

CDK11-cyclin L-SAP30BP grids were screened on a Glacios cryo-TEM operated at 200kV acceleration voltage and equipped with a Falcon 4i direct electron detector. Data were acquired using the EPU software (Thermo Fisher Scientific) in EER format at a pixel size of 0.5675 Å/pixel and a total exposure of 60 electrons/Å^2^, using aberration free image shift (AFIS) to accelerate the data collection. 500 or more movies, each fractionated into 40 frames for motion correction, were processed on the fly using cryoSPARC live (version 4.4.1) ^53^, enabling assessment of particle quality and orientation distribution by streaming 2D classification and streaming 3D refinement. Suitable grids were progressed to high-resolution data collection.

### High-resolution cryo-EM data collection

For high resolution structure determination, data were acquired on Titan Krios G2 and G3i cryo-TEMs operated at 300 kV acceleration voltage and equipped with a K3 camera and a BioQuantum energy filter (Gatan Inc.). Data were collected using the EPU software (versions 3.4 and 3.7; Thermo Fisher Scientific) in TIFF format at 165,000x nominal magnification using the hardware binning mode of the K3 camera, resulting in a pixel size of 0.504 Å/pixel (AMP-PNP complex) or 0.51 Å/pixel (OTS964-containing complexes). Movies were acquired using a defocus range of -0.8 µm to -1.8 µm and a total electron exposure of 70-72 electrons/Å^2^, fractionated into approx. 70 frames. Aberration-free image shift and fringe-free imaging were used for acceleration of the data collection. Grid squares were selected manually and brought to eucentric height using EPU (Thermo Fisher Scientific), before automated detection of holes and subsequent hole selection by grid-specific filtering of ice thickness. For the CDK11-cyclin L-SAP30BP complex bound to AMP-PNP, 16,936 and 19,396 exposures were acquired from two grids (referred to as AM4-3 and BG210-3 below); for the OTS964-bound CDK11-cyclin L-SAP30BP complex 13,829 exposures from one grid (AM10-3); for the CAK-OTS964 complex 10,788 exposures from one grid (BG61-1); and for the CDK2-cyclin A-OTS964 complex 15,468 exposures from one grid (BG65-1).

### Cryo-EM data processing

Data were initially processed in cryoSPARC live (version 4.4.1) ^53^ before transfer to RELION ^54^ (version 5.0-beta incorporating BLUSH regularisation ^55^) for additional processing and high-resolution 3D refinement (Supplementary Fig. 8). In cryoSPARC live, raw movies were motion corrected with 2x binning, resulting in pixel sizes of approx. 1 Å/pixel in the motion-corrected micrographs. Following inspection of CTF fit, relative ice thickness and max. per-frame motion, lower quality micrographs were rejected. Particle picking used circular and elliptical blob settings (80-110 Å diameter) to extract particles in a box size of 160 x 160 pixels before Fourier cropping to 80 x 80 pixels, resulting in approx. 2 Å pixel size in the extracted particles. Following streaming 2D classification and streaming 3D refinement during live processing, additional 2D classification to maximise retrieval of high-quality particles, particularly from rare views, was performed. Particles from 8-10 good classes obtained from streaming 2D classification during the cryoSPARC live session were re-classified into 50 2D-classes using a batch-size of 300 particles for 100 iterations. In parallel, all extracted particles also underwent re-classification into 100 2D-classes with a batch-size of 300 particles for 100 iterations. Particles of selected good classes from both re-classifications were combined and duplicate particles removed before re-extracting and re-centring of the final particle set. The metadata of the particles selected by this procedure (approx. 1,600,000 from AM4-3 and approx. 2,600,000 from BG210-4) were exported and converted to RELION-compatible *.star files using PyEM programs ^56^ before import into RELION 5.0.

In RELION 5.0, particles were re-extracted from the cryoSPARC motion-corrected micrographs using coordinate information contained in the imported particles. Extracted particles then underwent masked 3D auto-refinement using a previous reconstruction of the same complex from Glacios screening as an initial model and with BLUSH regularisation ^55^ enabled, reaching ∼2.5 Å resolution. Particles were then subjected to 3D classification with no alignment (regularisation parameter t=20) and 3D auto-refined again. For all but one dataset (CDK11-cyclin L-SAP30BP grid AM4-3), the remaining particle images were re-extracted at a larger box size (192 x 192 pixels) to prevent aliasing of the high resolutions in Fourier space. For the data from grid AM4-3, another 3D classification step was inserted before re-extraction to further improve the quality of the retained particles (Supplementary Fig. 8). Subsequently, CTF refinement (beam tilt and trefoil) was applied, followed by high-resolution 3D auto-refinement. For the CDK11-cyclin L-SAP30BP reconstruction in complex with AMP-PNP, data from two grids were merged at this point, leading to a further resolution increase. All other structures (i.e. the OTS964-bound complexes) were each determined from a single dataset collected on a single grid. For these datasets, CTF refinement was applied twice because of unexpectedly large beam tilt values that initially limited the obtained resolution.

Map resolution assessment by Fourier shell correlation (FSC), orientation distribution plots, and assessment of resolution anisotropy using the 3D FSC validation server 57 are provided in Supplementary Figs 2, 5, and 6.

### Model building and refinement

Atomic models were iteratively built in COOT ^58^ and refined using the real space refinement algorithm in PHENIX ^59^. Starting models for atomic model building were generated using AlphaFold3 ^30^ for CDK11-cyclin L-SAP30BP or from deposited structures of human CAK ^50^ and CDK2-cyclin A ^47^. Refinement restraints for the OTS964 ligand were generated in PHENIX ELBOW ^60^. Refined models were validated using MOLPROBITY ^61^ as implemented in PHENIX ^62^. Refinement statistics are provided in Supplementary Tables 2 and 3.

### Analysis and depiction of structures

Buried surface areas at protein-protein interfaces were computed using the PDBePISA web server at the European Bioinformatics Institute (http://www.ebi.ac.uk/pdbe/prot_int/pistart.html) ^31^. Figures depicting molecular complexes were generated using USCF Chimera X ^63^ and PyMOL (The PyMOL Molecular Graphics System, version 2.5.2-2.5.6, Schrodinger LLC).

## Data accessibility

The cryo-EM maps and atomic coordinate models for CDK11-cyclin L-SAP30BP-AMPNP, CDK11-cyclin L-SAP30BP-OTS964 (1 EMDB and 2 PDB entries), CAK-OTS964, and CDK2-cyclin A-OTS964 structures have been deposited in the EMDB with accession codes EMD-53224, EMD-53221, EMD-53205, and EMD-53204 and in the PDB with accession codes 9QKZ, 9QKT, 9QL1, 9QJN, and 9QJJ. Maps with reduced sharpening B-factor (B = - 10) used in some figures for visualisation of more dynamic protein segments have been supplied as additional maps within the EMDB entries of the CDK11B-cyclin L2-SAP30BP structures. Mass spectrometry data have been deposited to the ProteomeXchange Consortium via the PRIDE partner repository with accession code PXD060582.

## Acknowledgments

We thank Teige Matthews-Palmer for help with electron microscopy, Ruth Knight for help with insect cell culture, Christopher Richardson for help with high-performance computing, and Shruti Mittal, Caroline Ewens, and Rob van Montfort for support of biophysical assays. We thank Vlad Pena, Max Douglas, Jonathon Pines, and Ioana Maruntel for discussions. We acknowledge the Addgene plasmid repository and depositors John Chodera, Nicholas Levinson, and Markus Seeliger, from whom we obtained the human CDK2 plasmid, and Jonathon Pines, from whom we obtained the human cyclin A sequence. B.J.G. was supported by a career development fellowship from the Medical Research Council of the UK (MR/V009354/1) and A.J.S.M. was funded by an ICR PhD studentship. We acknowledge Diamond and our local contact Éilís Bragginton for access and support of the cryo-EM facilities at the UK national electron Bio-Imaging Centre (eBIC), proposal BI33974 (session BI33974-9), and we acknowledge data collection at the London Consortium for High-Resolution Cryo-EM (LonCEM) facility, supported by Wellcome Grant No. 206175/Z/17/Z and partner institutes.

## Author contributions

A.J.S.M. cloned, expressed, and purified CDK11-containing protein complexes. V.I.C. prepared CDK2-cyclin A and CAK. A.J.S.M. and B.J.G. prepared cryo-EM specimens and performed initial cryo-EM screening. A.J.S.M., N.B.C., and B.J.G. acquired cryo-EM data. A.J.S.M. and B.J.G. processed cryo-EM data. B.J.G. and A.J.S.M. built and refined the atomic models. A.J.S.M. performed kinase assays and biochemical experiments supported by S.J.H.. T.I.R. and J.C. performed and oversaw mass spectrometry analysis. C.A. developed and provided the cryo-EM peptide. B.J.G. and A.J.S.M. performed structure interpretation and drafted the initial version of the manuscript; all authors contributed to its final form. B.J.G. designed and oversaw the project.

