## Supplementary Information for "Cryo-EM structures of the CDK11-cyclin L-SAP30BP complex reveal mechanisms of CDK11 regulation"

**This file includes:**
Supplementary Figures 1-8
Supplementary Tables 1-3

**Additional supplementary materials include:**
Supplementary Dataset 1

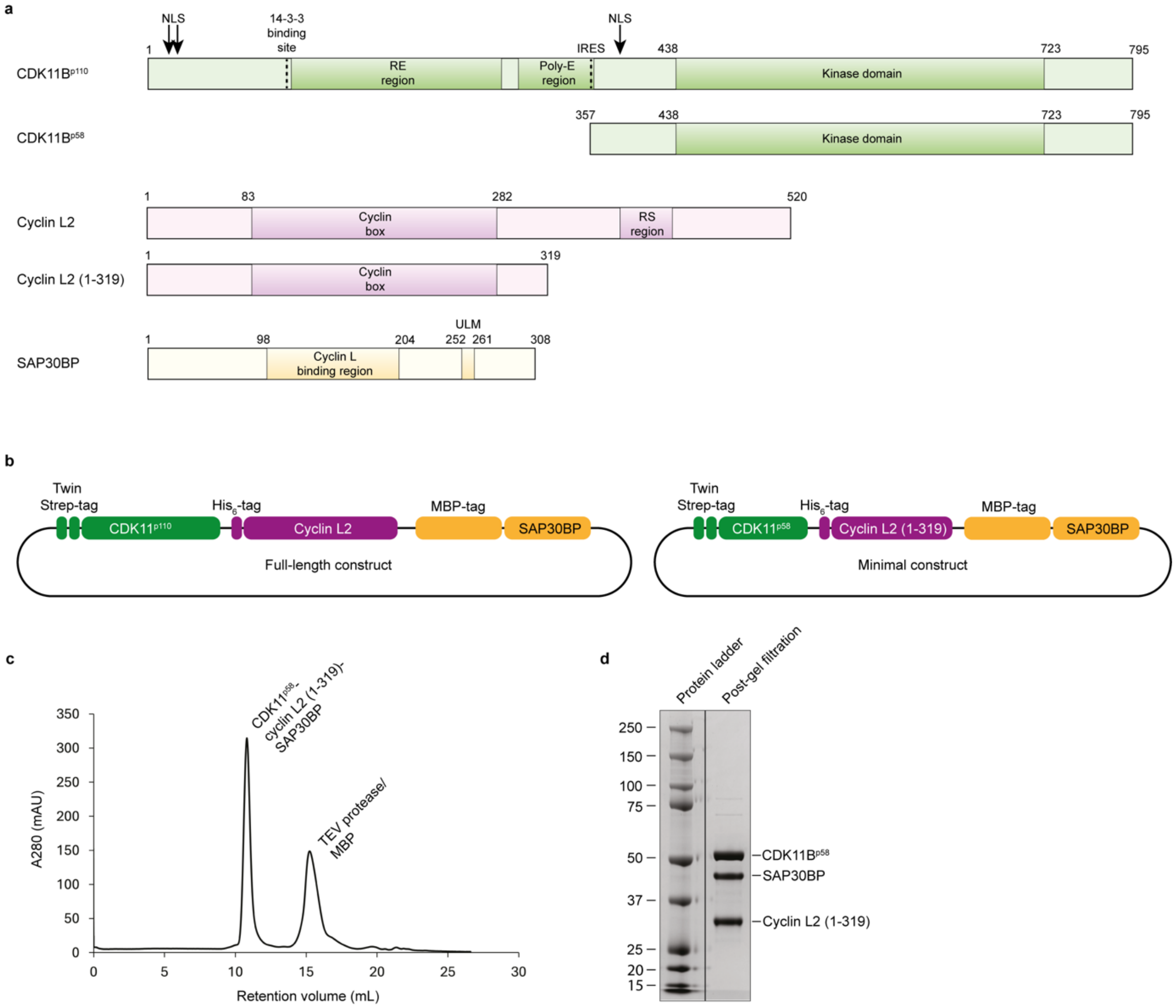


**Supplementary Figure 1 | Expression and purification of the CDK11-cyclin L-SAP30BP complex.** (**a**) Domain organisation of CDK11B, cyclin L2, and SAP30BP. Protein domains, important binding sites, and notable low-complexity regions are indicated. Abbreviations: NLS, nuclear localisation signal; RE, arginine-glutamate-rich region; RS, arginine-serine-rich region; IRES, internal ribosome entry site; ULM, U2AF-ligand motif. (**b**) Expression constructs for production of full length (CDK11B$^{p110}$, cyclin L2, SAP30BP) and minimal (CDK11B$^{p58}$, cyclin L2 (1-319), SAP30BP) CDK11-cyclin L-SAP30BP complexes. (**c**) Size exclusion chromatogram from purification of the minimal construct. (**d**) Purified protein preparation of the minimal construct (used for structure determination).

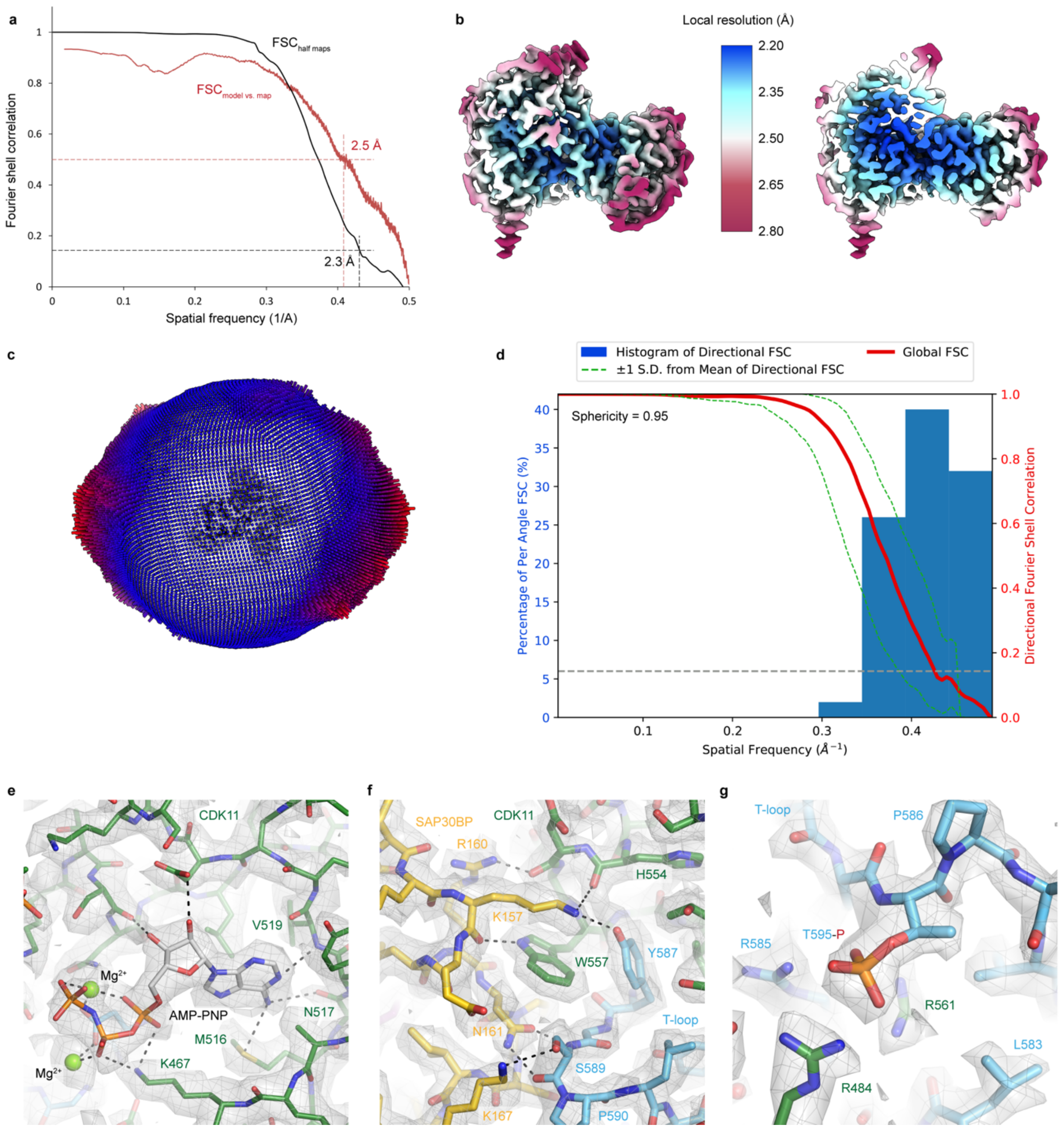


**Supplementary Figure 2 | Validation and quality of the CDK11-cyclin L-SAP30BP cryo-EM reconstruction.** (**a**) Half-map and model vs. map resolution estimates for the CDK11-cyclin L-SAP30BP structure at FSC = 0.143 and FSC = 0.5, respectively [64]. (**b**) Local resolution estimation for the CDK11-cyclin L-SAP30BP structure. (**c**) Orientation distribution plot of the CDK11-cyclin L-SAP30BP cryo-EM reconstruction. (**d**) Analysis of the CDK11-cyclin L-SAP30BP cryo-EM reconstructions by 3D FSC [57]. (**e**, **f**) Sections of the cryo-EM map of the CDK11-cyclin L-SAP30BP complex. Density is shown in semi-transparent grey surface and mesh. CDK11 is shown in green (T-loop cyan) and SAP30BP in yellow. (**g**) Molecular environment of phosphorylated CDK11 T595 within the T-loop (cyan) with surrounding arginine residues.

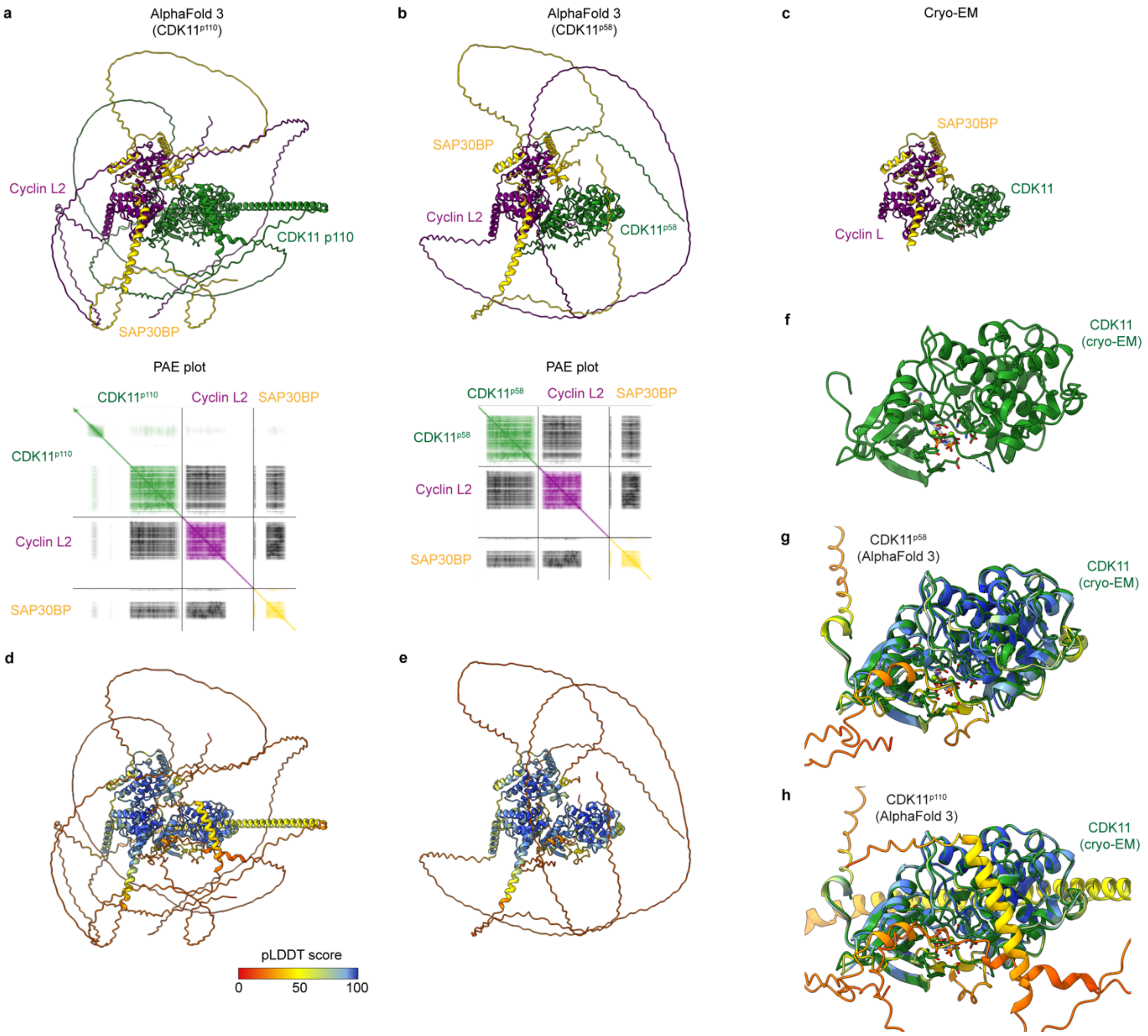


**Supplementary Figure 3 | CDK11$^{p110}$ and CDK11$^{p58}$ AlphaFold 3 predictions and comparison to the cryo-EM structure.** (**a**) Predicted structure of the CDK11B$^{p110}$-cyclin L2-SAP30BP complex (top) and plot of the predicted aligned error (bottom). (**b**) Predicted structure of the CDK11B$^{p58}$-cyclin L2-SAP30BP complex (top) and plot of the predicted aligned error (bottom). (**c**) Cryo-EM structure of the CDK11B-cyclin L2-SAP30BP complex. (**d**, **e**) Plots of the pLDDT score for the structure prediction of the CDK11$^{p110}$- and CDK11$^{p58}$-containing complexes. Segments not resolved in the cryo-EM structure mostly carry low pLDDT scores and are likely poorly ordered or flexibly attached in solution. (**f**-**h**) Superposition of the CDK11B structure derived from our cryo-EM data (green, f) with the predictions of the CDK11B$^{p58}$ and CDK11B$^{p110}$ structures (g, h, coloured by pLDDT score). The extensions not observed in our cryo-EM structures carry low pLDDT scores and are likely disordered in the free complex.

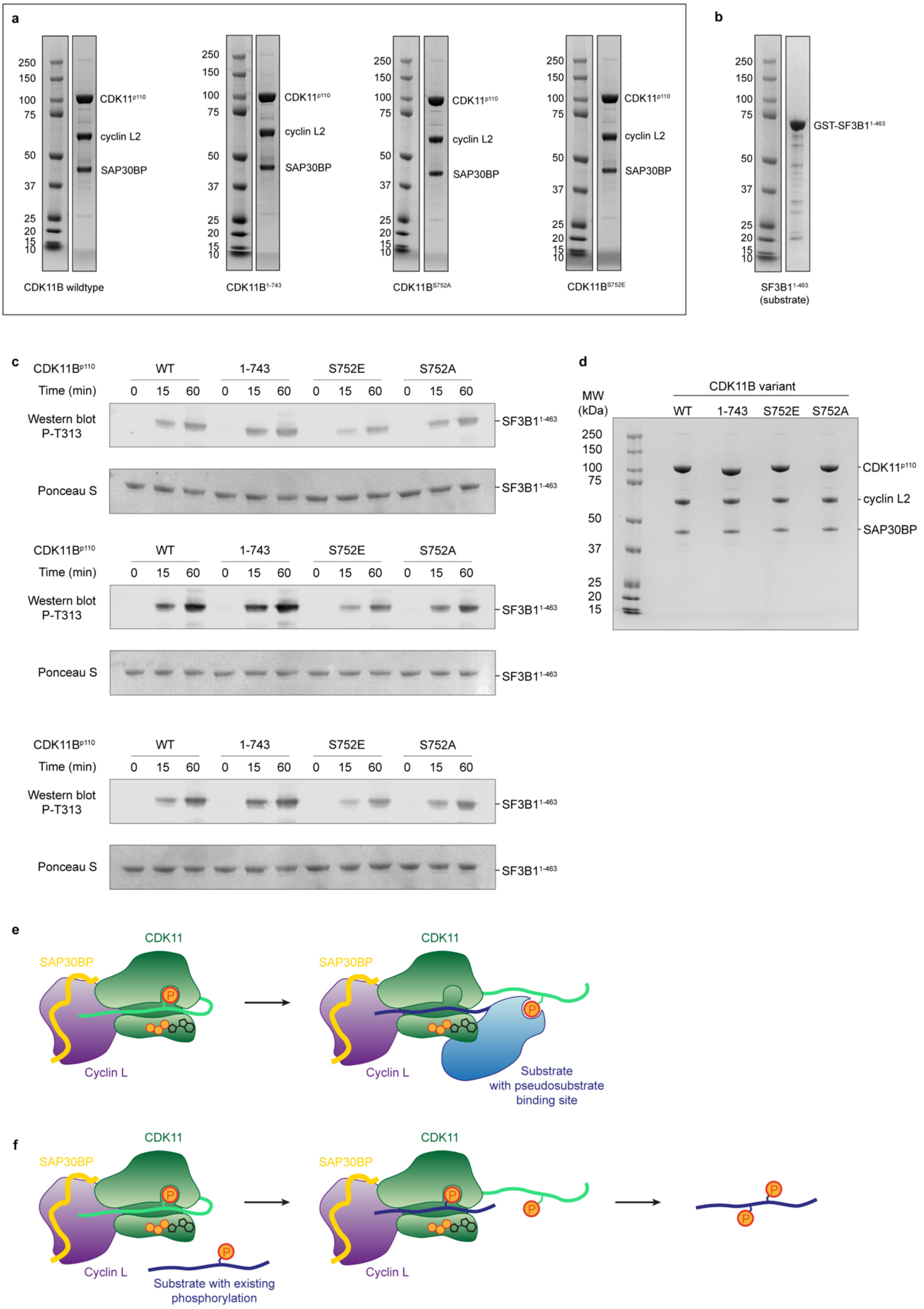


**Supplementary Figure 4 | Purification of proteins and protein complexes for kinase assays.** (**a**) Coomassie-stained SDS-PAGE gel lanes that show purified complexes containing mutant variants of CDK11B$^{p110}$ used for kinase assays. (**b**) Purification of the assay substrate GST-SF3B1$^{1-463}$. (**c**) Western blots and Ponceau S stained membranes of three additional replicates of kinase assays used for statistics in Fig. 3g. (**d**) Kinase input control to verify that calculated dilutions of CDK11 complexes used in kinase assays resulted in equivalent amounts of kinase complex. (**e**) Schematic representation of a hypothesis that may reconcile our finding of reduced CDK11 activity in the presence of a S752 phosphorylation mimic with the previously documented role of S752 phosphorylation in stimulating CDK11 activity towards certain targets. The substrate binding site is schematically represented as a cleft, and the active site is indicated by a stylised ATP molecule. (**f**) Schematic representation of CDK11 binding of a substrate with pre-existing phosphorylation adjacent to a CDK11 target site.

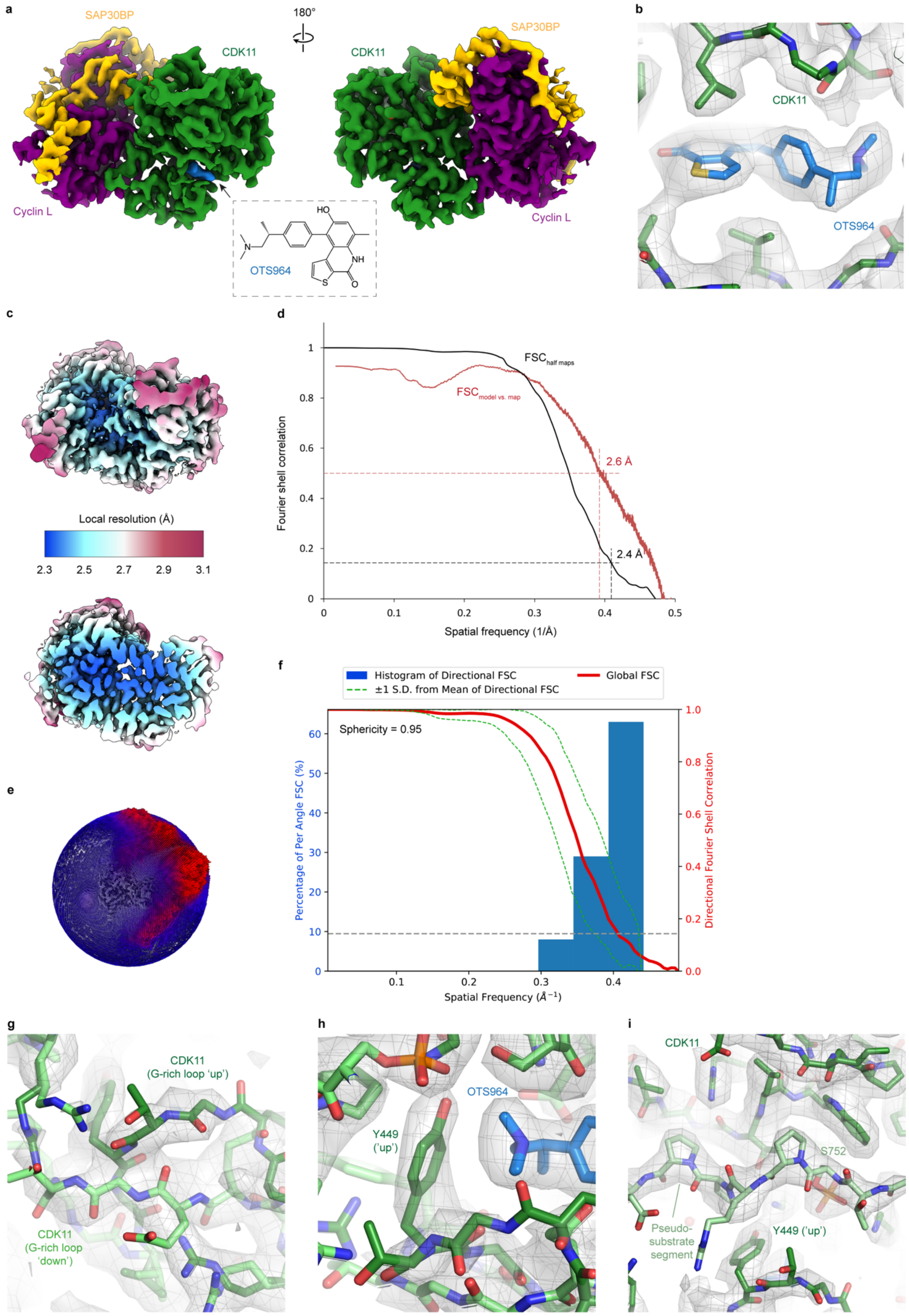


**Supplementary Figure 5 | 3D reconstruction and validation of the CDK11-cyclin L-SAP30BP-OTS964 complex.** (**a**) Depiction of the CDK11-cyclin L-SAP30BP-OTS964 cryo-

EM structure and the chemical structure of OTS964. CDK11 is shown in green, cyclin L in purple, SAP30BP in yellow, and OTS964 in blue. (**b**) Rendering of OTS964 (blue) in the cryo-EM map of the CDK11-cyclin L-SAP30BP-OTS964 complex (shown as semi-transparent grey mesh and surface). (**c**) Local resolution estimation for the CDK11-cyclin L-SAP30BP-OTS964 structure. (**d**) Half-map and model vs. map resolution estimates for the CDK11-cyclin L-SAP30BP-OTS964 structure at FSC = 0.143 and FSC = 0.5, respectively [64]. (**e**) Orientation distribution plot of the CDK11-cyclin L-SAP30BP-OTS964 cryo-EM reconstruction. (**f**) Analysis of the CDK11-cyclin L-SAP30BP-OTS964 cryo-EM reconstruction by 3D FSC [57]. (**g**, **h**) Two conformations of the CDK11 G-rich loop (shown in dark green and lime) with the cryo-EM density shown as semi-transparent grey mesh and surface. The conformations are shown in superposition because we were unable to separate the two G-rich loop conformations computationally. (**i**) The CDK11 pseudo-substrate segment (light green; phosphorylated S752 is labelled) shown in the cryo-EM map of the OTS964-bound CDK11-cyclin L-SAP30BP complex.

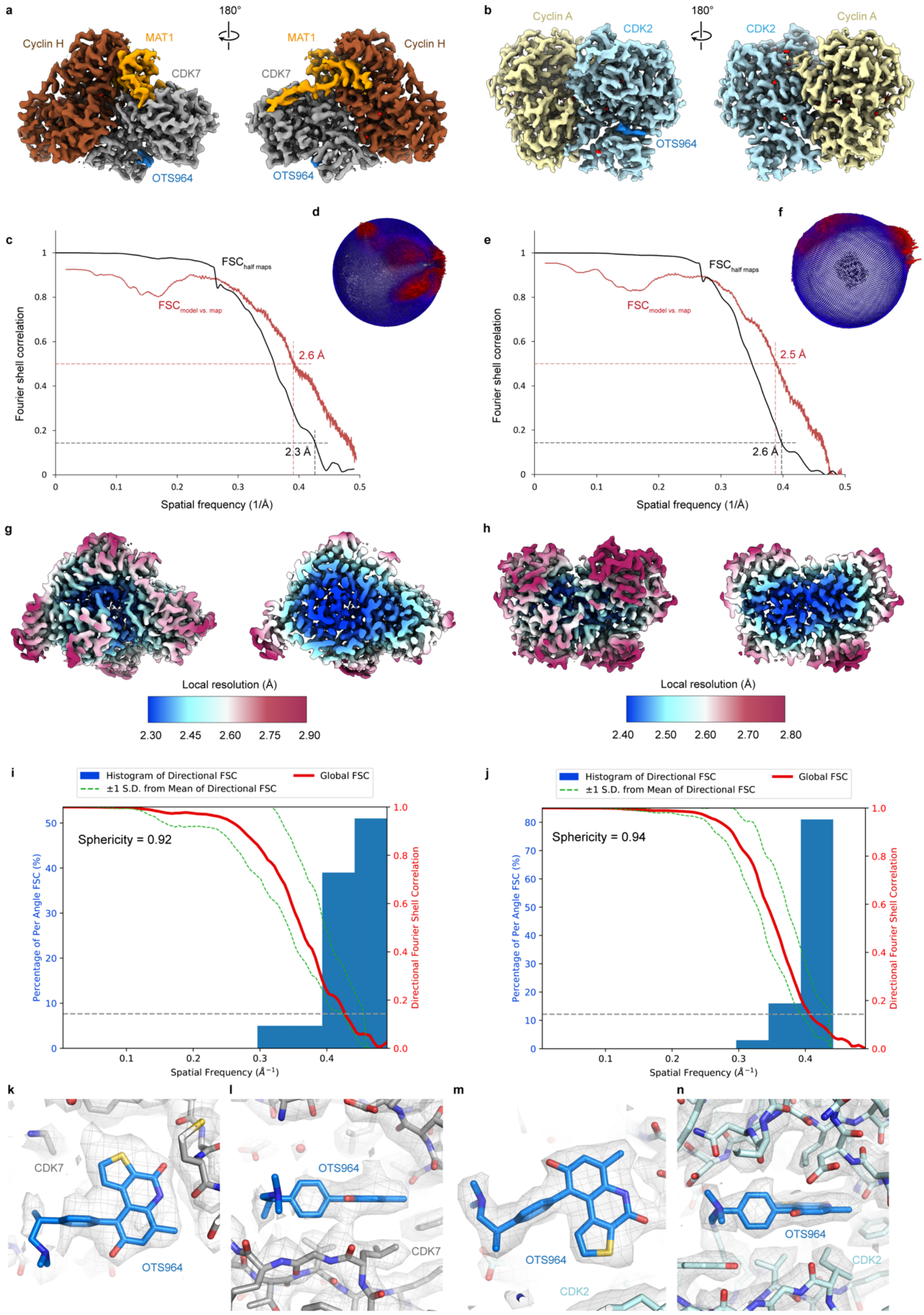


**Supplementary Figure 6 | 3D reconstruction and validation of CAK-OTS964 and CDK2-cyclin A-OTS964 complexes.** (**a**) Cryo-EM map of CAK-OTS964 at 2.3 Å resolution. CDK7 is shown in grey, MAT1 in orange, cyclin H in brown, and OTS964 in blue. (**b**) Cryo-EM map of CDK2-cyclin A-OTS964 at 2.5 Å resolution. CDK2 is shown in cyan, cyclin A in light yellow, and OTS964 in blue. (**c**) Half-map and model vs. map resolution estimates for the CAK-OTS964 structure at FSC = 0.143 and FSC = 0.5, respectively [64]. (**d**) Orientation distribution of the CAK-OTS064 cryo-EM reconstruction. (**e**, **f**) As c, d, but for CDK2-cyclin A-OTS964. (**g**, **h**) Local resolution estimation for the CAK-OTS964 (g) and CDK2-cyclin A-OTS964 (h) cryo-EM reconstructions. (**i**, **j**) Analysis of the CAK-OTS964 (i) and CDK2-cyclin A-OTS964 (j) cryo-EM reconstructions by 3D FSC [57]. (**k**, **l**) Two views of OTS964 (blue) in the cryo-EM map of the CAK-OTS964 complex (CDK7 shown in light grey, cryo-EM density shown as a semi-transparent grey mesh and surface). (**m**, **n**) Two views of OTS964 (blue) in the cryo-EM map of the CDK2-cyclin A-OTS964 complex (CDK7 shown in light cyan, cryo-EM density shown as a semi-transparent grey mesh and surface).

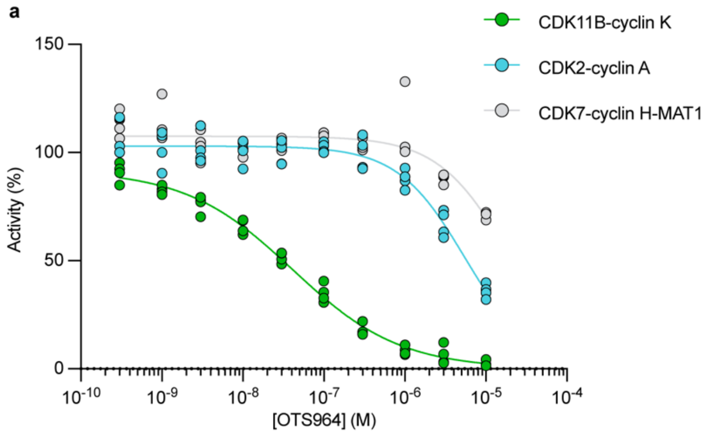


b

| Kinase | $IC_{50}$ (nM) |
|---|---|
| CDK11-cyclin K | 41 (30, 54) |
| CDK2-cyclin A | 5,500 (4500, 6700) |
| CDK7-cyclin H-MAT1 | >10,000 (N. D.) |

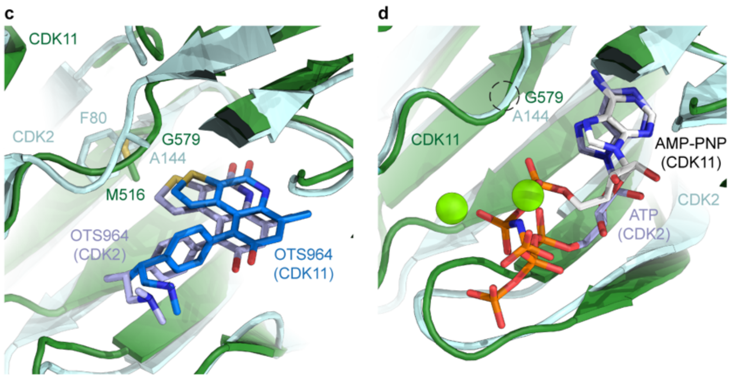


**Supplementary Figure 7. Verification of relative affinity of OTS964 to CDK11-, CDK2-, and CDK7-containing complexes and analysis of inhibitor selectivity.** (**a**) Enzyme inhibition assay to determine the inhibitory properties of OTS964 against CDK7-cyclin H-MAT1 and to compare performance of the OTS964 sample used for structural studies against literature data for CDK11B and CDK2 [28]. N = 4 data points (technical replicates) were recorded per condition (inhibitor concentration and kinase). (**b**) $IC_{50}$ values derived from the data shown in a. Borders of confidence intervals (95%) are shown in brackets. The results indicate that the OTS964 sample used for cryo-EM experiments performs in line with prior data for CDK11B and CDK2 [28]. OTS964 is even more inefficient in inhibiting CDK7 than it is for CDK2. (**c**) Comparison between CDK11-cyclin L-SAP30BP-OTS964 and CDK2-cyclin A-OTS964 highlighting the different size of the gatekeeper residues (M516 and F80, respectively) and the structural difference at CDK11 and CDK2 residues G579 and A144, respectively. (**d**) Unlike in the inhibitor-bound state, the backbone conformation around CDK11 G579/CDK2 A144 is almost identical in the nucleotide-bound state of the two kinases (CDK2-cyclin A: PDB ID 1FIN).

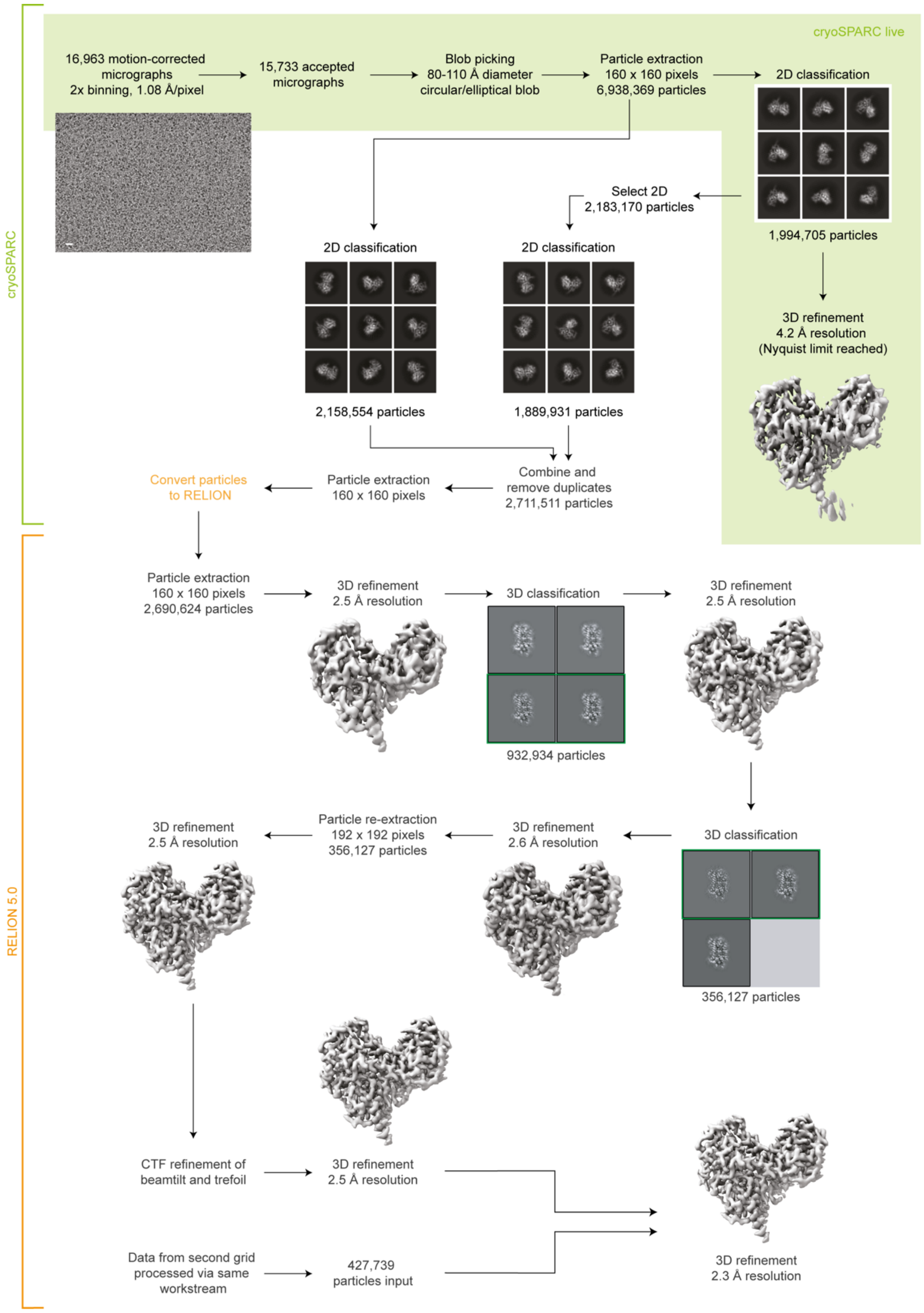


**Supplementary Figure 8 | Data processing workflow for structure determination of the CDK11-cyclin L-SAP30BP complex.** Data from two grids were processed independently using the outlined workflow. Merging of 356,127 particle images from the first grid and 427,739 particle images from the second grid yielded the final reconstruction at 2.3 Å resolution. Scale bar for sample micrograph: 100 Å.

**Supplementary Table 1 | Summary of mass spectrometry analysis of phosphorylated residues in recombinant CDK11-cyclin L-SAP30BP.** Protein: Name of the protein. Only data for CDK11B, cyclin L2, and SAP30BP are shown; detected peptides for low-abundance contaminants are included in Supplementary Dataset 1. Residue: Phosphorylatable residues within detected phosphopeptides. Probability: Probability of a phosphate being located on a residue, given its position in the sequence in case of multiple possible phospho-acceptors in a peptide. Density: Evidence of phosphate in the cryo-EM density. Phosphates observed with high probability in mass spectrometry but not in the cryo-EM map may be present sub-stoichiometrically in our sample.

| Protein | Residue | Probability (%) | Density |
|---|---|---|---|
| CDK11B | S384 | 98.7 | Residue not modelled |
| | T395 | 97.5 | Residue not modelled |
| | S398 | 33 | Residue not modelled |
| | Y406 | 33 | Residue not modelled |
| | S410 | 100 | Residue not modelled |
| | S414 | 100 | Residue not modelled |
| | S589 | 100 | Phosphate not observed |
| | T595 | 100 | Phosphate visualised |
| | Y615 | 25 | Phosphate not observed |
| | S616 | 99.2 | Phosphate not observed |
| | T617 | 25 | Phosphate not observed |
| | S623 | 25 | Phosphate not observed |
| | T726 | 100 | Phosphate not observed |
| | T751 | 99.6 | Phosphate not observed |
| | S752 | 100 | Phosphate visualised |
| Cyclin L2 | S43 | 100 | Residue not modelled |
| | T62 | 100 | Residue not modelled |
| | S239 | 100 | Phosphate not observed |
| SAP30BP | S9 | 98.9 | Residue not modelled |
| | Y14 | 100 | Residue not modelled |
| | S18 | 100 | Residue not modelled |
| | S22 | 100 | Residue not modelled |
| | S33 | 100 | Residue not modelled |
| | S104 | 100 | Phosphate not observed |
| | S106 | 99.7 | Phosphate not observed |
| | S113 | 100 | Phosphate not observed |
| | S255 | 99.1 | Residue not modelled |
| | S259 | 33.2 | Residue not modelled |
| | T264 | 33.2 | Residue not modelled |
| | T265 | 33.2 | Residue not modelled |

**Supplementary Table 2 | Data collection and refinement statistics, part 1.**

| **Dataset (kinase / ligand)** | **CDK11 / AMP-PNP** | **CDK11 / OTS964** | |
|---|---|---|---|
| Microscope | Titan Krios G2 | Titan Krios G3i | |
| Stage type | Autoloader | Autoloader | |
| Voltage (kV) | 300 | 300 | |
| Detector | Gatan K3 | Gatan K3 | |
| Energy filter | Bio Quantum | Bio Quantum | |
| Acquisition mode | 2x hardware binning | 2x hardware binning | |
| Pixel size (non-superresolution) (Å) | 0.504 | 0.51 | |
| Defocus range (μm) | 0.8-1.8 | 0.8-1.8 | |
| Electron exposure ($e^-/A^2$) | 70 | 70 | |
| **Reconstruction** | **EMD-53224** | **EMD-53221** | |
| Software | RELION 5.0 beta | RELION 5.0 beta | |
| Particles after 2D classification | 3,610,927 + 2,690,624 | 2,243,141 | |
| Particles final | 783,863 | 286,108 | |
| Box size (pixels) | 192 x 192 x 192 | 192 x 192 x 192 | |
| Final pixel size (Å) | 1.008 | 1.02 | |
| Accuracy rotations (°) | 1.36 | 1.26 | |
| Accuracy translations (Å) | 0.42 | 0.40 | |
| Map resolution (Å) | 2.3 | 2.4 | |
| Map resolution range (Å) | 2.2-3.4 | 2.3-3.5 | |
| Sphericity | 0.95 | 0.95 | |
| Map sharpening B-factor ($Å^2$) | -61.5 | -53.7 | |
| **Coordinate refinement** | | | |
| Software and algorithm | PHENIX (real space refine) | PHENIX (real space refine) | |
| Resolution cutoff (Å) | 2.3 | 2.4 | |
| $FSC_{model-vs-map}$=0.5 (Å) | 2.5 | 2.6 | |
| **Model** | **PDB-9QKZ** | **PDB-9QKT** | **PDB-9QL1** |
| Number of residues | 728 | 762 | 762 |
| Protein | 680 | 681 | 681 |
| Ligand (AMP-PNP / OTS964 / $Mg^{2+}$ / $H_2O$) | 1 / 0 / 2 / 45 | 0 / 1 / 0 / 80 | 0 / 1 / 0 / 80 |
| B-factors overall | 84.1 | 74.8 | 74.4 |
| Protein | 84.2 | 75.1 | 74.7 |
| Ligand (AMP-PNP, OTS964, $Mg^{2+}$ / $H_2O$) | 85.3 / 72.5 | 53.1 / 66.0 | 55.7 / 65.8 |
| R.M.S. deviations | | | |
| Bond lengths (Å) | 0.003 | 0.04 | 0.04 |
| Bond angles (°) | 0.47 | 1.39 | 1.40 |
| **Validation** | | | |
| Molprobity score | 1.56 | 1.71 | 1.64 |
| Molprobity clashscore | 7.14 | 5.98 | 5.54 |
| Rotamer outliers (%) | 1.64 | 3.28 | 2.79 |
| $C_\beta$ deviations (%) | 0.0 | 0.0 | 0.0 |
| Ramachandran plot | | | |
| Favored (%) | 98.2 | 98.7 | 98.3 |
| Allowed (%) | 1.6 | 1.1 | 1.5 |
| Outliers (%) | 0.2 | 0.2 | 0.2 |
| Rama-Z scores | | | |
| Whole | 2.55 | 2.64 | 2.74 |
| Helix | 2.42 | 2.66 | 2.71 |
| Sheet | -0.60 | 0.03 | 0.64 |
| Loop | 1.23 | 0.96 | 0.99 |

**Supplementary Table 3 | Data collection and refinement statistics, part 2.**

| **Dataset** | **CDK2 / OTS964** | **CDK7 / OTS964** |
|---|---|---|
| Microscope | Titan Krios G3i | Titan Krios G3i |
| Stage type | Autoloader | Autoloader |
| Voltage (kV) | 300 | 300 |
| Detector | Gatan K3 | Gatan K3 |
| Energy filter | Bio Quantum | Bio Quantum |
| Acquisition mode | 2x hardware binning | 2x hardware binning |
| Pixel size (non-superresolution) (Å) | 0.51 | 0.51 |
| Defocus range (μm) | 0.6-1.6 | 0.6-1.8 |
| Electron exposure ($e^-/A^2$) | 70 | 70 |
| **Reconstruction** | **EMD-53204** | **EMD-53205** |
| Software | RELION 5.0 beta | RELION 5.0 beta |
| Particles after 2D classification | 3,482,979 | 2,312,870 |
| Particles final | 503,775 | 340,642 |
| Extraction box size (pixels) | 192 x 192 x 192 | 192 x 192 x 192 |
| Final pixel size (Å) | 1.02 | 1.02 |
| Accuracy rotations (°) | 1.48 | 1.21 |
| Accuracy translations (Å) | 0.42 | 0.40 |
| Map resolution (Å) | 2.5 | 2.4 |
| Map resolution range (Å) | 2.4-3.3 | 2.2-3.4 |
| Sphericity | 0.94 | 0.92 |
| Map sharpening B-factor ($Å^2$) | -69 | -27 |
| **Coordinate refinement** | | |
| Software and algorithm | PHENIX (real space refine) | PHENIX (real space refine) |
| Resolution cutoff (Å) | 2.5 | 2.4 |
| $FSC_{model-vs-map}$=0.5 (Å) | 2.6 | 2.6 |
| **Model** | **PDB-9QJJ** | **PDB-9QJN** |
| Number of residues | 602 | 773 |
| Protein | 546 | 630 |
| Ligand (AMP-PNP / OTS964 / $Mg^{2+}$ / $H_2O$) | 0 / 1 / 0 / 55 | 0 / 1 / 0 / 142 |
| B-factors overall | 37.0 | 62.0 |
| Protein | 37.2 | 62.0 |
| Ligand (AMP-PNP, OTS964, $Mg^{2+}$ / $H_2O$) | 23.1 / 30.5 | 90.0 / 56.6 |
| R.M.S. deviations | | |
| Bond lengths (Å) | 0.04 | 0.04 |
| Bond angles (°) | 1.55 | 1.48 |
| **Validation** | | |
| Molprobity score | 1.35 | 1.32 |
| Molprobity clashscore | 4.49 | 5.39 |
| Rotamer outliers (%) | 1.4 | 1.1 |
| $C_\beta$ deviations (%) | 0.0 | 0.0 |
| Ramachandran plot | | |
| Favored (%) | 99.1 | 98.4 |
| Allowed (%) | 0.9 | 1.4 |
| Outliers (%) | 0.0 | 0.2 |
| Rama-Z scores | | |
| Whole | 1.68 | 1.73 |
| Helix | 1.75 | 2.39 |
| Sheet | 0.42 | -0.65 |
| Loop | 0.61 | -0.29 |
